# Systematic analysis of membrane contact sites in *Saccharomyces cerevisiae* uncovers modulators of cellular lipid distribution

**DOI:** 10.1101/2021.10.17.464712

**Authors:** Inês Gomes Castro, Shawn P. Shortill, Samantha K. Dziurdzik, Angela Cadou, Suriakarthiga Ganesan, Emma Joanne Fenech, Hadar Meyer, Amir Fadel, Yotam David, Michael Davey, Carsten Mattes, Rosario Valenti, Robert Ernst, Vanina Zaremberg, Tim P. Levine, Christopher J. Stefan, Elizabeth Conibear, Maya Schuldiner

## Abstract

Actively maintained close appositions, or contact sites, between organelle membranes, enable the efficient transfer of biomolecules between the various cellular compartments. Several such sites have been described together with their tethering machinery. Despite these advances we are still far from a comprehensive understanding of the function and regulation of most contact sites. To systematically characterize the proteome of contact sites and support the discovery of new tethers and functional molecules, we established a high throughput screening approach in *Saccharomyces cerevisiae* based on co-localization imaging. We imaged split fluorescence reporters for six different contact sites, two of which have never been studied before, on the background of 1165 strains expressing a mCherry-tagged yeast protein that have a cellular punctate distribution (a hallmark of contact sites). By scoring both co-localization events and effects on reporter size and abundance, we discovered over 100 new potential contact site residents and effectors in yeast. Focusing on several of the newly identified residents, we identified one set of hits as previously unrecognized homologs to Vps13 and Atg2. These proteins share their lipid transport domain, thus expanding this family of lipid transporters. Analysis of another candidate, Ypr097w, which we now call Lec1 (<u>L</u>ipid-droplet <u>E</u>rgosterol <u>C</u>ortex 1), revealed that this previously uncharacterized protein dynamically shifts between lipid droplets and the cell cortex, and plays a role in regulation of ergosterol distribution in the cell.

## Introduction

The hallmark of eukaryotic cells is the presence of organelles as biochemically distinct compartments. To effectively coordinate cellular responses, organelles must communicate and work cooperatively. Membrane contact sites play an essential role in this communication by actively tethering two organelles in proximity to each other thus enabling a direct physical interaction and the exchange of ions, lipids and other small molecules (Eisenberg-Bord et al., 2016).

Contact sites are formed between most, if not all, cellular membranes (Kakimoto et al., 2018; Shai et al., 2018; Valm et al., 2017) and are implicated in a growing number of processes from organelle morphology and inheritance, to lipid metabolism and intracellular signaling (Prinz et al., 2020). To perform these functions, contact sites harbor a defined membrane composition, enriched with specific proteins and lipids that form tethers between the opposing membranes, enabling a functional interaction (Scorrano et al., 2019). While several contact sites have been observed for decades and we know of multiple tethers and functions for the few well-studied contacts, the formation and function of the majority of contact sites remains poorly understood. One important step towards such an understanding would be to obtain the complete proteome of contact site residents.

Several techniques have been developed and adapted to enable the identification of contact site proteins, including proximity labeling assays and systematic high throughput screening approaches (Cho et al., 2017; Shai et al., 2018; van Vliet et al., 2017). Although these approaches generally require previous knowledge of contact site resident proteins, this can be circumvented by the use of synthetic reporters, such as split proteins, targeted to the opposing membranes of two organelles. When in close proximity, these split proteins will interact and tether both organelles, either emitting a fluorescent signal (Cieri et al., 2018; Eisenberg-Bord et al., 2016; Yang et al., 2018) or performing an enzymatic reaction (Cho et al., 2020; Kwak et al., 2020).

We have previously taken advantage of the split fluorescence protein approach to identify unknown contact sites, and to uncover two new tethering proteins for the peroxisome-mitochondria (PerMit) contact site in *Saccharomyces cerevisiae* (from here on termed yeast) (Shai et al., 2018). Here, we build on this approach and expand it by using high content screens to systematically analyze effectors and resident proteins of six different contact sites: PerMit, Lipid Droplet-Endoplasmic Reticulum (LD-ER) contact (LiDER), nuclear ER-vacuole junction (NVJ), peroxisome-vacuole contact (PerVale), Plasma Membrane (PM, or cortex)-LD contact (pCLIP) and Golgi-peroxisome contact (GoPo), the latter two not having been studied before. All together, we have identified 158 unique proteins with a potential role in tethering and/or regulation of these 6 contact sites. While focusing on the pCLIP, we identified 3 proteins – Fmp27 and Ypr117w (renamed Hob1 and Hob2 respectively, for <u>Hob</u>bit homologs 1 and 2), and Csf1, which share homology with the lipid transporters Vps13 and Atg2, suggesting that other members of this family of proteins are localized to contact sites. Additionally, as we explored the LiDER, we identified a protein of unknown function, Ypr097w. We demonstrated that Ypr097w plays a role in regulating the distribution of ergosterol in yeast cells, and suggest to name it Lec1 for <u>L</u>ipid-droplet <u>E</u>rgosterol <u>C</u>ortex 1. Collectively, our studies highlight the power of our high content screening approach for discovering novel functions of uncharacterized proteins and for expanding our understanding of contact site biology as well as better grasping how cells actively distribute cellular lipids.

## Results

### Mapping the proteome of multiple contact sites using high throughput microscopy screens uncovers potential residents and regulators

To identify new potential contact site resident proteins and regulators we used a set of contact site reporters based on a bimolecular fluorescence complementation assay (Alford et al., 2012; Sung and Huh, 2007) that we and others have previously used to visualize contact sites (Cieri et al., 2018; Kakimoto et al., 2018; Shai et al., 2018; Tashiro et al., 2020). In short, each part of a split-Venus protein is used to tag membrane proteins on opposing organelles. Only in instances where these organelles come into extremely close proximity, as is the case at contact sites, the two parts complement each other to form a full fluorophore. Since any membrane protein with the correct topology (terminus facing the cytosol) can be used for this assay (Shai et al., 2018), this approach allows for the visualization of contact sites in the absence of known components. Taking advantage of this we used the previously verified reporters for the PerMit, NVJ, LiDER, PerVale and pCLIP (Shai et al., 2018) and a new reporter for the GoPo (Fig 1 S1A). The NVJ reporter served as a positive control, as it is one of the more characterized contacts, therefore in an effective screen we would expect to identify many of its known components. The remaining contacts are considerably less well described and presented a new challenge.

To identify potential new contact site proteins, we generated strains expressing each of the above-mentioned reporters and crossed them against a collection of strains overexpressing mCherry-tagged proteins (Fig. 1A). This collection is a subset of the full genome SWAT *TEF2pr*-*mCherry* library (Weill et al., 2018; Yofe et al., 2016), in which every yeast protein is tagged at its N-terminus with mCherry and overexpressed under the *TEF2* promoter. This subset contains 1165 strains where the mCherry-tagged protein is either localized to intracellular puncta – the typical cellular distribution of contact site proteins – or has previously been identified as a contact site protein (Table S1). These strains were mated against each of the 6 reporter strains using an automated mating, sporulation and haploid selection procedure (Cohen and Schuldiner, 2011; Tong and Boone, 2006), to create six new collections where each haploid strain expresses a split-Venus reporter and overexpresses one mCherry-tagged protein (Fig.1A). Each collection was subsequently imaged by high throughput microscopy and the resulting images were manually analyzed for full or partial co-localization between the contact site reporter and the mCherry-tagged protein (residents), and/or effect of the overexpressed protein on the contact site reporter number and/or brightness (effectors). All strains classified as hits were isolated and re-imaged on a higher resolution spinning disk confocal microscope (Micro 1; see Materials and Methods). Only those strains that were still clear hits after this step were included in the final hit list (Table S2). It is important to note that in the case of small punctate organelles (such as LDs, peroxisomes and Golgi) the resolution of light-microscopy does not enable us to differentiate between co- localization to the contact or distribution over the entire organelle. Hence, if a known resident protein of such organelles appeared as co-localizing with the reporter in the screen, we did not label it as a “contact resident” and only considered it a hit in the screen if it was also an “effector”.

**Figure 1.**
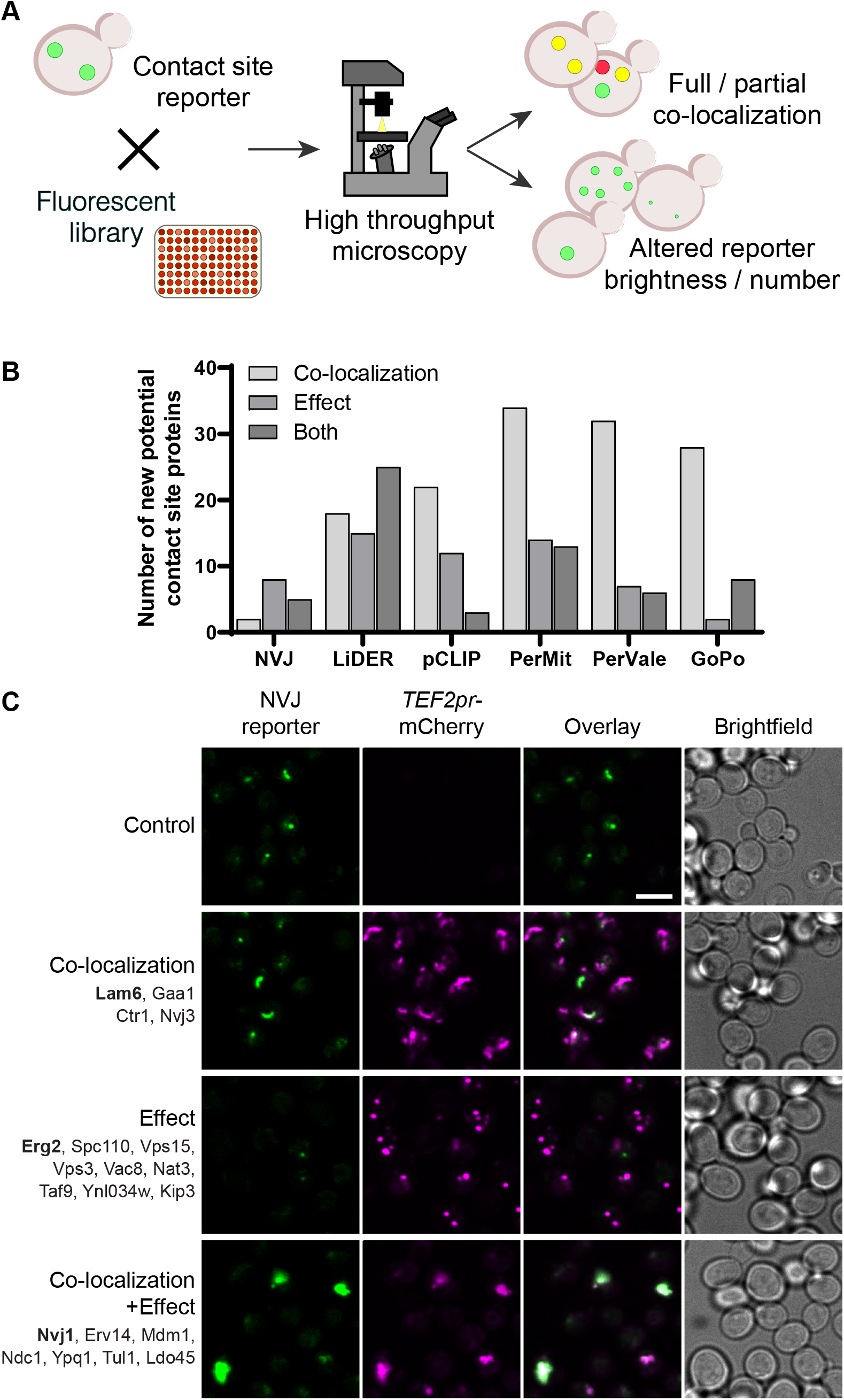
Systematic screens uncover new contact site residents and effectors in yeast. (**A**) Schematic representation of the high throughput screening approach used to identify new contact site residents and regulators. (**B**) Graphical representation of the number of newly suggested contact site residents and effectors (excluding known contact site residents), categorized based on co-localization with- and/or effect on contact site reporter. (**C**) Summary results of the NVJ contact site screen. A total of 20 proteins were identified and characterized based on their co-localization with and/or effect on the contact site reporter. Individual hits are listed under each category, and example images are from proteins highlighted in bold. Scale bar, 5 µm; Images obtained using Micro 1 (For details of each microscope used see “Materials and methods” section).

A total of 158 unique proteins were identified across all six contact sites (Table S2, Fig. 1B). When looking at our positive control, the NVJ (Fig. 1C), we found 20 proteins that co-localize with and/or affect the contact. It was reassuring to see that known residents, such as Nvj1, Nvj3, Mdm1 and Lam6 (Elbaz-Alon et al., 2015; Henne et al., 2015; Murley et al., 2015; Pan et al., 2000), all co- localized with the contact site reporter. Additionally, overexpression of mCherry-Erg2 reduced the frequency and brightness of the reporter signal, suggesting that modulation of the ergosterol biosynthesis pathway affects the formation of the NVJ. This is particularly interesting, since several NVJ resident proteins play a role in sterol sensing and transport, specifically Lam5, Lam6 and Osh1 (Gatta et al., 2015; Levine and Munro, 2001; Murley et al., 2015), and this contact has also been shown to play a role in the mevalonate pathway by facilitating the assembly of HMG-CoA reductases, and consequently affecting activity of these enzymes, during acute glucose restriction (Rogers et al., 2021). This result supports the ability of our screen to capture not only putative tethers but also effectors.

### The pCLIP contact reveals a family of proteins structurally related to Vps13

One interesting yet uncharacterized contact site in yeast is the pCLIP, between the PM and LDs, where we identified 37 proteins that co-localize with and/or affect the contact (Fig. 2A). To understand how such proteins can function in the contact, we first focused on proteins with known roles in contact sites or with domains that are commonly found in contact site proteins, such as lipid-binding/transfer domains. From these we identified Mdm1, a known contact site protein that localizes to a three-way LD-ER-vacuole contact (Hariri et al., 2019). Interestingly, the *Drosophila melanogaster* homolog of Mdm1, Snazarus, also localizes to a triple contact that instead involves LD-ER-PM membranes (Ugrankar et al., 2019). Although Mdm1 and Snazarus bind LDs via different domains, and have different specificity for membrane lipids, this new localization for Mdm1 suggests it could also play a role at a yeast LD-ER-PM contact site.

**Figure 2.**
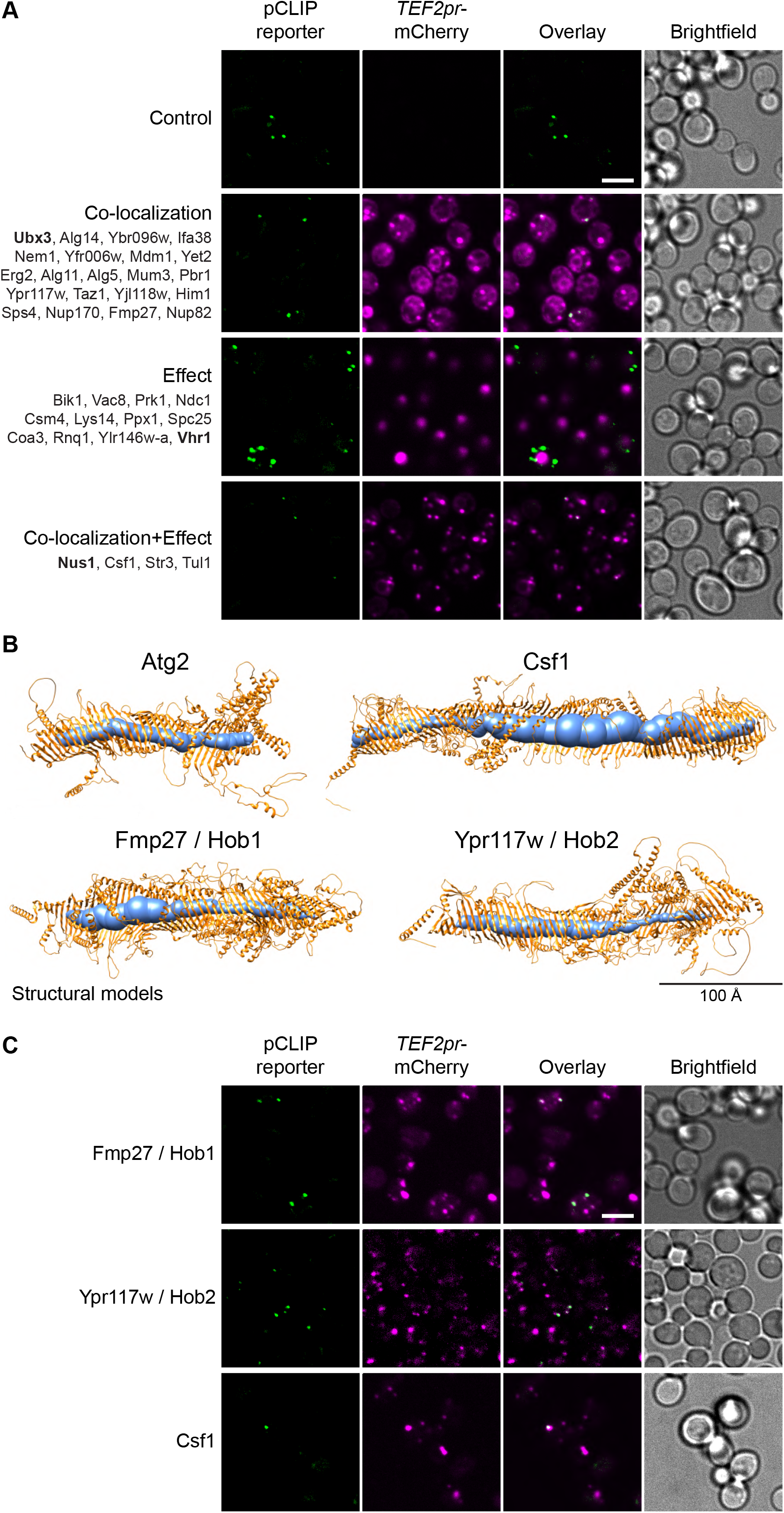
The LD-PM screen reveals new Vps13 homologs. (**A**) Summary results of the LD-PM contact site screen. A total of 37 proteins were identified and characterized based on their co-localization with and/or effect on the contact site reporter. Individual hits are listed under each category, and example images are from proteins highlighted in bold. Scale bar, 5 µm; Images obtained using Micro 1. (**B**) Predicted structures of Hob1, Hob2 and Csf1 in comparison to Atg2. Structural predictions were created by the AlphaFold2 consortium for the full-length proteins Hob1/2 and Atg2. For Csf1, four overlapping regions were predicted separately using the AlphaFold2 Colab (Mirdita et al., 2021). Internal channels were modelled with MOLE and are depicted in blue. (**C**) N-terminally mCherry-tagged Hob1, Hob2 and Csf1, under the strong *TEF2* promoter, co- localize with the pCLIP contact site reporter. Scale bar, 5 µm; Images obtained using Micro 1.

Additionally, we identified a set of 3 poorly characterized proteins – Fmp27, Ypr117w (closely related paralogs that we now name Hob1 and Hob2, respectively, for <u>Hob</u>bit homologs 1 and 2, due to their homology to the *D. melanogaster* Hobbit protein) and Csf1. We found that these are structural homologs of AsmA, a bacterial relative of Vps13 (Fig. 2 S1A). Advanced modelling indicates that they all share full-length homology to Vps13 and Atg2 (Fig. 2B, Fig. 2 S1B), two well described long, tubular lipid transfer proteins (Kumar et al., 2018; Li et al., 2020; Maeda et al., 2019; Osawa et al., 2019; Valverde et al., 2019). In yeast, Vps13 has been shown to reside to both the NVJ and the mitochondria-vacuole contact (vCLAMP) (Lang et al., 2015; Park et al., 2016). Like Vps13 and Atg2, these proteins are predicted to be made up almost entirely of long, predominantly hydrophobic channels, which can potentially transport multiple lipid molecules (Fig. 2B, Fig. 2 S2A). C-terminally tagged versions of Hob1, Hob2 and Csf1 mainly localize to ER-PM contact sites (Neuman et al., 2022; Peter et al., 2021). Hob1 and Hob2, and their *Drosophila* homologue Hobbit, target contact sites through N-terminal transmembrane domains that anchor in the ER (Fig. 2 S2B), and C-terminal elements that bind other organelles including the PM (Neuman et al., 2022). Csf1 also has a predicted N-terminal ER anchor (Fig. 2 S2B) and has been shown to play a role in lipid homeostasis (Peter et al., 2021). When tagged at the N-terminus, which may interfere with ER targeting, all 3 proteins co-localize with pCLIP, suggesting that these proteins function at this contact site (Fig. 2C). Csf1 overexpression appears to decrease the number of detectable pCLIP foci and increase the number of LiDER contacts, indicating a possible crosstalk between these sites (Table S2). However, further analysis of this protein is needed to characterize these effects. Moreover, the identification of Hob1, Hob2 and/or Csf1 at several different contacts (LiDER, pCLIP, PerMit) suggests that these proteins, like yeast Vps13 and human VPS13A, VPS13C and ATG2A (Bean et al., 2018; Kumar et al., 2018; Tamura et al., 2017; Tang et al., 2019; Yeshaw et al., 2019), function at multiple contacts.

### The LiDER contact screen uncovers a previously uncharacterized protein

One contact that has been little studied is the ER-LD contact site or LiDER. Since LDs remain in close connection to the ER throughout their life cycle in yeast (Hugenroth and Bohnert, 2020) this contact is quite prevalent. In our screen the LiDER reporter co-localized with and/or was affected by 59 proteins (Fig. 3A). Similar to the pCLIP, we focused our analysis on proteins with domains that are common to contact site proteins, such as lipid-binding/transfer domains. As expected, we found the known LiDER protein Mdm1 amongst the hits for this screen. Additionally, we identified Ypr097w, a protein of unknown function that both co-localized with and increased the brightness and frequency of the LiDER reporter in cells (Fig. 3A, bottom panel). Ypr097w has a Phox homology (PX) domain, which is present in Mdm1 and a large number of proteins conserved across all eukaryotic kingdoms, and which interacts with several anionic lipids, including the phosphoinositide species phosphatidylinositol 3-phosphate (PI3P) (Chandra et al., 2019; Yu and Lemmon, 2001). Additionally, Ypr097w contains a predicted FFAT motif (two phenylalanines, FF, in an acidic tract), a conserved amino acid sequence that enables the interaction of several proteins with the major sperm protein (MSP) domain of VAMP-associated protein (VAP) proteins (Scs2 and Scs22 in yeast) (Slee and Levine, 2019). This sequence is present in several contact site proteins, where it enables the formation of contact sites between the ER and other organelles (Murphy and Levine, 2016). These characteristics suggest that Ypr097w could function at the LiDER contact.

**Figure 3.**
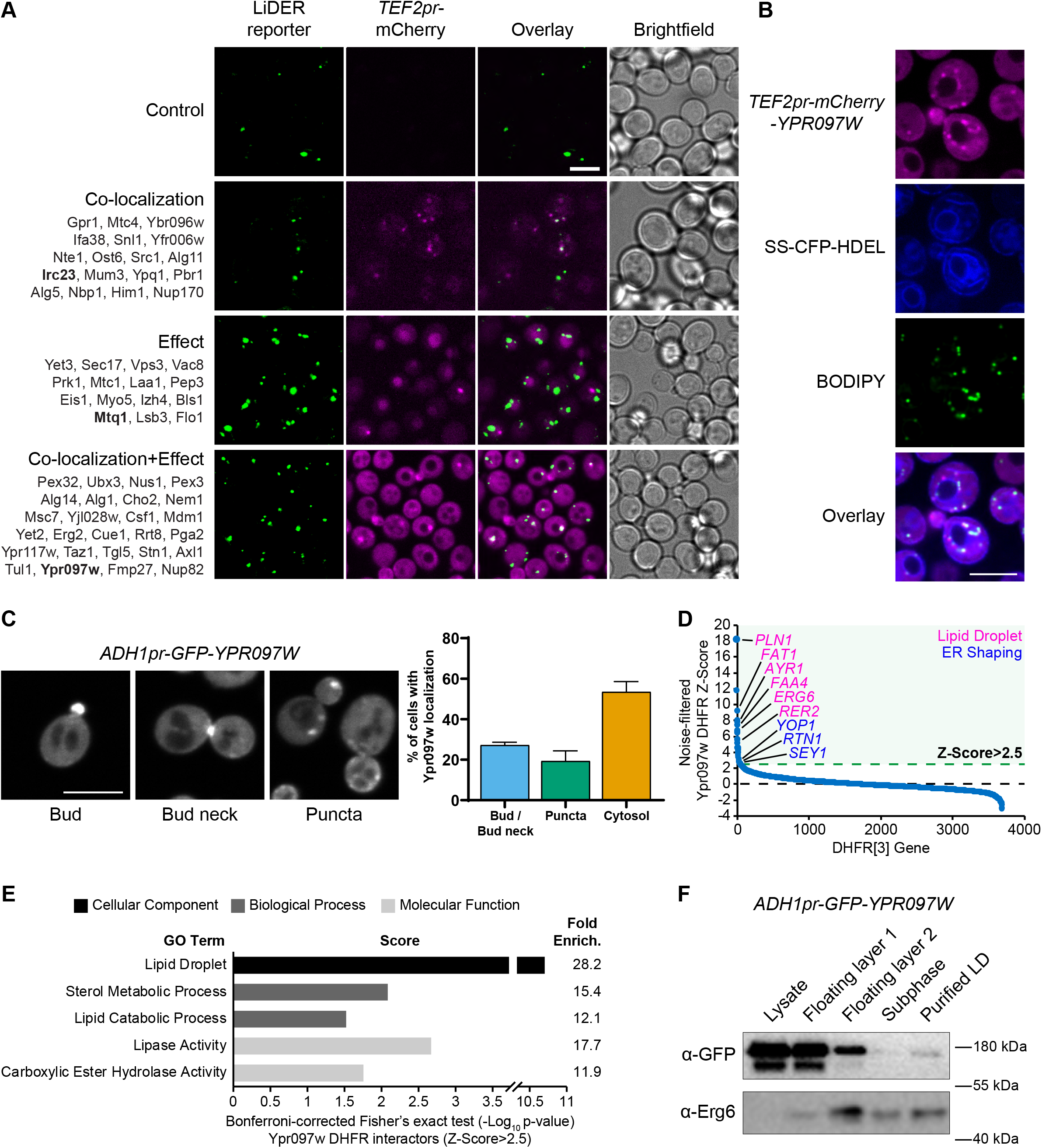
The ER-LD screen reveals new potential lipid-binding proteins. (**A**) Summary results of the LiDER contact site screen. A total of 59 proteins were identified and characterized based on their co-localization with and/or effect on the contact site reporter. Individual hits are listed under each category, and example images are from proteins highlighted in bold. Scale bar, 5 µm; Images obtained using Micro 1. (**B**) mCherry-Ypr097w, under control of the *TEF2* promoter, localizes to LDs (BODIPY) in the periphery of the ER (SS-CFP-HDEL). Scale bar, 5 µm; Images obtained using Micro 2. (**C**) Cellular localization of *ADH1pr-GFP-YPR097W* during mid-logarithmic growth. Cells were categorized based on Ypr097w localization and quantified as shown in the graph. Scale bar, 5 µm; Images obtained using Micro 1. (**D**) Noise-filtered Z-Score distribution of colony area from the DHFR protein fragment complementation assay with endogenously expressed Ypr097w used as a bait. Filtering was used to remove prey strains that exhibited strong signal with a validated cytoplasmic DHFR reporter. An enrichment of proteins with reported LD subcellular localization patterns and proteins with roles in ER shaping was observed in prey strains with a Z-Score of >2.5. (**E**) Functional enrichment analysis of strong Ypr097w DHFR interactors (Z>2.5) using the Gene Ontology enrichment analysis tool (Ashburner et al., 2000; Mi et al., 2019; The Gene Ontology Consortium, 2019). GO terms are presented as the negative base 10 log of the associated p-value from a Bonferroni-corrected Fisher’s exact test. Lipid Droplet (GO:0005811); Sterol Metabolic Process (GO:0016125); Lipid Catabolic Process (GO:0016042); Lipase Activity (GO:0016298); Carboxylic Ester Hydrolase Activity (GO:0052689) are significantly enriched ontologies. (**F**) Cells expressing *ADH1pr-GFP-YPR097W* were collected and fractionated by centrifugation to obtain enriched LD fractions. Sequential fractions were run by SDS-page and analyzed by western blot. GFP-Ypr097w is present in the LD fraction. Erg6 was used as a LD marker.

To confirm the localization of Ypr097w in the absence of a synthetic reporter we re-tagged it at the N-terminus with mCherry under the control of a strong promoter (Fig. 3B) and imaged its location relative to both the ER and LDs. In these conditions, Ypr097w localized to internal puncta, several of which co-localize with the LD stain, BODIPY, on the interface with the ER (Fig. 3B), supporting our original observation.

When using the endogenous promoter to tag Ypr097w with GFP at either the N- or C-terminus, Ypr097w showed a strikingly different localization than when overexpressed – it was found primarily at buds and bud necks, with relatively few internal puncta (Fig. 3 S1A). Tagging at either termini maintains a functional Ypr097w, as will be described below in Fig. 5A. Because endogenous levels were difficult to detect, we also imaged cells expressing GFP-Ypr097w from the moderate constitutive *ADH1* promoter (Fig. 3C). In this strain, GFP-Ypr097w was more clearly detected and showed a similar distribution pattern to the endogenous promoter, localizing to the bud and bud neck with some increase in the number of internal puncta (Fig. 3C). Additionally, many cells (nearly 60%) were devoid of a punctate signal and appeared to have a general cytosolic localization. This suggests that the overexpression of Ypr097w either alters cellular physiology or saturates binding sites on the bud/bud neck and causes intracellular accumulation.

As an alternative approach to identify the cellular environment occupied by Ypr097w under endogenous expression levels, and to reconcile these diverse observations, we performed a genome-wide protein fragment complementation screen based on a split dihydrofolate reductase (DHFR) enzyme (Tarassov et al., 2008) (Table S3). This screen, which measures the interactions by growth as colonies over multiple days, showed that the strongest interactors (Z-Score>2.5), which represent proteins in close proximity to Ypr097w, consist primarily of well-characterized LD-localized enzymes (Fig. 3D-E) and include ER membrane proteins that localize to, and help shape, regions of tubular ER, such as Rtn1, Yop1 and Sey1 (Craene et al., 2006; Hu et al., 2009; Voeltz et al., 2006). In contrast, bud neck-localized proteins were not enriched in this screen. This suggests that under the nutrient conditions experienced in a yeast colony (as compared to cells grown in liquid media and during mid-logarithmic growth), Ypr097w is primarily localized to LDs in the proximity of the ER.

These distinct distribution patterns suggest that the localization of Ypr097w is dynamically regulated under different environmental conditions. Indeed, we found that when cells were shifted from nutrient-containing media to PBS, both endogenous (Fig. 3 S1B) and *ADH1pr*-expressed (not shown), GFP-Ypr097w rapidly (< 2 min) re-localized from bud/bud neck to the cytosol, an effect that was blocked if the PBS was supplemented with glucose. Similarly, longer glucose starvation in synthetic media led to a mostly cytosolic localization of this protein (Fig. 3 S1C). This change in localization could be a response to cytoplasmic acidification due to glucose depletion, and resulting loss of interaction of Ypr097w with pH sensing membrane lipids (Shin et al., 2020). In contrast, in stationary cells, GFP-Ypr097w localized mostly in a punctate pattern (Fig. 3 S1D, left panel), which quickly shifted to a mix of punctate and bud/bud neck upon replenishment with fresh media (Fig. 3 S1D, right panel). The movement of Ypr097w between buds and bud necks to internal puncta (many of which co-localize with LDs) under different growth conditions, suggests that Ypr097w acts as a nutrient sensor and may have a role at LDs primarily in lipid storage conditions.

To further examine the pool of Ypr097w associated with LDs, we biochemically purified LDs from cells expressing GFP-Ypr097w under control of the moderate constitutive *ADH1* promoter. We found that GFP-Ypr097w is present in purified LD fractions (Fig. 3F), and that it migrates as a doublet in cell lysates, suggesting that Ypr097w is post-translationally modified. In fact, several phosphorylation sites have been identified in Ypr097w in high throughput studies (Beltrao et al., 2012; Lanz et al., 2021), including phosphorylation of residue S451 in the region flanking the predicted FFAT motif (Fig. 3 S2A), which has the potential to regulate interaction with VAP proteins (Kumagai et al., 2014; Mattia et al., 2020). Moreover, the punctate localization of Ypr097w increased in cells lacking both yeast VAP proteins *(Δscs2/Δscs22*) (Fig 3 S2B). However, we found that expression of either phosphomimetic (S451E; S451D) or phospho-null (S451A) mutants (Fig. 3 S2C), or a mutant lacking the FFAT motif (Y465A,D458A; data not shown), in a strain lacking endogenous Ypr097w, did not show any clear differences in localization relative to the wild type (WT) Ypr097w control. This suggests that neither phosphorylation of S451 nor the presence of the FFAT motif regulates YPR097w targeting. The effect of *Δscs2/22* on Ypr097w localization could be mediated by redundant FFAT motifs in Ypr097w, through interactions with other FFAT-containing proteins, or by interactions with other surfaces of the VAP proteins.

Taken together, these results suggest that the localization of Ypr097w is regulated in response to environmental conditions, and that this protein dynamically partitions between the bud/bud neck and LDs. However, what drives this redistribution is not known and will form the basis of future studies.

### The predicted structure of Ypr097w reveals a large hydrophobic cavity and a new family of lipid binding proteins

To understand how Ypr097w is targeted to different cellular locations, we examined its predicted domain architecture. In addition to its PX domain, which harbors the predicted FFAT motif, Ypr097w has an annotated PX-associated domain (PXB) and a domain of unknown function (DUF) 3818 (Fig. 3 S2A) (Mistry et al., 2021). By testing a series of protein fragments, we found that truncations that perturb either the N or C-termini disrupt protein localization, and we did not identify any domain that was sufficient for localization to either LDs or to the bud/bud neck (Fig. 3 S2D).

Our finding that targeting is affected by truncation of either terminus of Ypr097w, both of which have no known structure, led us to analyze this protein using structural bioinformatics. Structural homology modeling by HHpred (Gabler et al., 2020) showed that the unnamed regions bracketing DUF3818 (Fig. 4A) have similar levels of conservation as the named domains, and that all regions, except the PX domain (which is longer than previously noted) are made up of multiple alpha-helices (data not shown). Together with our truncation results, this suggests that full length Ypr097w functions as a single unit. However, other than the PX domain, HHpred searches did not find any homology to solved structures or remote homology to other proteins. This implies that Ypr097w forms a previously undocumented fold.

**Figure 4.**
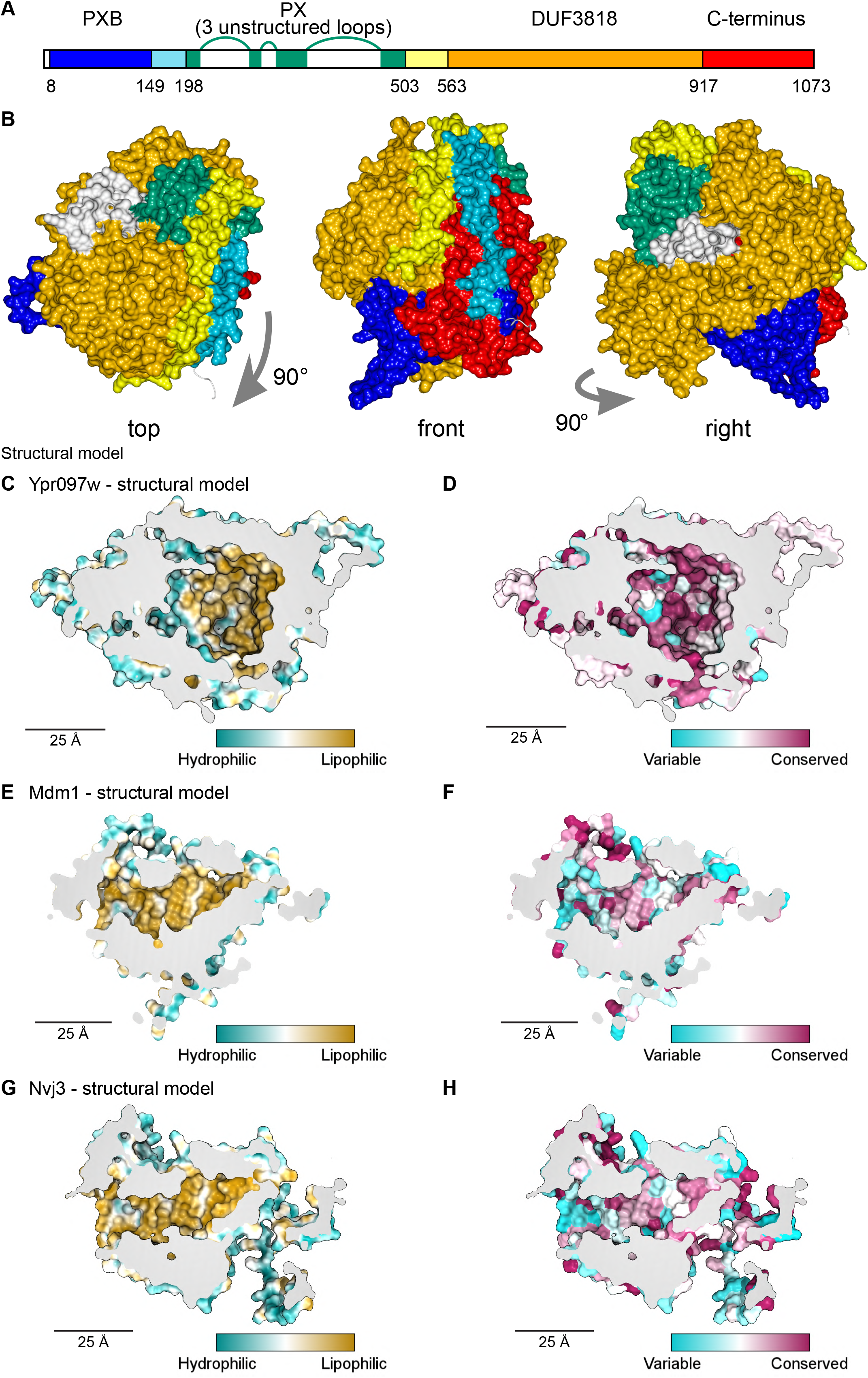
Structural bioinformatics predicts that Ypr097w forms a large spherical hydrophobic cavity shared with Mdm1 and Nvj3. (**A**) Domain map of Ypr097w based results from HHpred in which we could identify strand-1 of the PX domain (green) in residues 213-212, and find that the extreme C-terminus (red) is as helical and as conserved as the preceding DUF3818 domain (orange). The PX domain contains three loops without strong structural predictions (white). (**B**) AlphaFold2 structure prediction for Ypr097w, colored by domain as in (A), omitting loops outside the protein core. Three views from different angles. Domains are intimately associated with each other, in particular the PXB (blue) and extreme C-terminus (red). **(C-D)** Representations of the cavity of Ypr097w predicted by AlphaFold2, with the lining colored by either hydrophobicity (C) or conservation (D). **(E-F)** Representations of the cavity of Mdm1 predicted by AlphaFold2, coloring as for C/D respectively. **(G-H)** Representations of the cavity of Nvj3 predicted by AlphaFold2, coloring as for C/D respectively.

To determine the Ypr097w fold, we used *in silico* approaches, in particular AlphaFold2 (Jumper et al., 2021). AlphaFold2 outputs for Ypr097w and its full-length homologs in both *C. albicans* and *S. pombe* all predict that the different all-helical regions, which appear separate in the one-dimensional map (Fig. 4A), interact *in cis* to form a sphere ∼7 nm in diameter (Fig. 4B). A similar sphere is also predicted for a shorter *S. pombe* paralog that has PXB and DUF3818 without a PX domain (Fig. 4 S1A). A key aspect of these structures is the multiple inter-relationships between the different components of the primary structure. In particular the N-terminal PXB domain folds intimately together with the unnamed C-terminal region (Fig. 4B and Fig. 4 S1A). The PX domain is oriented to expose its PI3P binding site, which would allow Ypr097w to dock onto membranes (Fig. 4 S1B). Among the five AlphaFold2 models, the PX-negative *S. pombe* homolog is the most confidently predicted (Fig. 4 S1C), implying that the PX domain is unnecessary for the overall fold.

A striking feature of the predicted Ypr097w sphere is that it is hollow with an internal cavity (Fig. 4C-D). The surface of the cavity is mainly hydrophobic (Fig. 4C), a feature that is highly conserved (Fig. 4D), being present in all 5 modelled homologs (data not shown). Multiple tunnels potentially provide access from the cytosol (Fig. 4 S1D-F, arrows). Intimate interactions of N- and C-termini, overall spheroidal shape, a large internal hydrophobic-lined cavity, and tunnels linking the exterior to the interior were also predicted by the trRosetta server (data not shown) (Yang et al., 2020). Among the five proteins modelled by AlphaFold2, *S. cerevisiae* Ypr097w is unique in having three disordered loop inserts in its PX domain (Fig. 4A). Part of the first loop was modelled as dipping into the cavity (Fig. 4 S1D, modelled in white). However, since this loop is only found in the budding yeast protein, is predicted with low confidence (Fig. 4 S1E) and is not supported by other structural prediction algorithms and under certain running parameters is predicted to form part of the sphere wall (Fig. 4 S1F) we calculated the internal cavity without the loop and found that it has a volume of 29 nm^3^ (Fig. 4 S1C), equivalent to a sphere diameter of 3.9 nm.

Given the remarkable structure of Ypr097w, and the now readily available structures predicted by AlphaFold2, we decided to use a candidate approach to look for additional spherical domains with large lipophilic cavities among known contact site proteins in yeast, including our >100 hits. We noted that the contact site protein Mdm1, similarly to Ypr097w, has a PX domain and two associated domains of unknown structure – the PXA and the nexin-C domains – which are found only in Mdm1 and its homologs, including Nvj3 which like Mdm1 also has PXA domain and nexin- C domain (identified here using HHpred) at its N and C-termini. Strikingly, AlphaFold2 analysis of these proteins revealed that the PXA and nexin-C domains in Mdm1 and Nvj3 fold together to form a large spherical domain with an interior hydrophobic surface (Fig. 4E and G) that is highly conserved (Fig. 4F and H). While the PXA/nexin-C domains share no detectable sequence homology to Ypr097w, the structural similarities suggest that these proteins may share a lipid storage or transfer function.

### Ypr097w affects the cellular distribution of ergosterol

Since Ypr097w is localized to LDs, we assayed whether it has any effect on this organelle and its ability to store lipids. To test this, we first looked at the number of LDs in WT, *Δypr097w* and overexpressed *YPR097W* (under control of the *TDH3pr)* cells using the LD marker BODIPY. However, we observed no clear changes in the number of LDs (Fig. 5 S1A) or LD morphology (Fig. 5 S1B).

To test for changes in the total levels of storage lipids, we performed thin layer chromatography (TLC) of WT, *Δypr097w* and *TDH3pr-YPR097W* during mid-logarithmic growth, in stationary cultures (24h and 48h), and following growth resumption during new media replenishment (Fig. 5 S1C). When compared to control cells, no clear changes in the levels of diacylglycerol (DAG), triacylglycerol (TAG), ergosterol (Erg) or sterol esters (SE) were observed in any of the conditions tested. Additionally, lipidomic analysis of the overexpression strain relative to a control strain also did not uncover any major global lipid differences (Fig. 5 S1D, Table S4). Small significant differences were detected in the levels of CDP-DAG during mid-logarithmic phase and Lyso Phosphatidic Acid (LPA) in stationary cells (Fig. 5 S1D, insert), but these should be considered with caution, as the overall levels of these lipid classes in the samples were very low (up to 32pmol per sample) (Table S4). Overall, this suggests that that Ypr097w does not affect the global cellular levels of tested lipids.

While the overall lipid levels were not altered, we wondered if their cellular distribution may be. To test this we assayed the sensitivity of these strains to the drug Amphotericin B (AmB), which has been shown to bind accessible ergosterol at the PM (Gray et al., 2012), and could therefore reveal changes in cellular levels and distribution of free ergosterol. Surprisingly, despite the lack of changes in overall ergosterol or SE levels observed by TLC and lipidomics, loss of Ypr097w sensitized cells to AmB (Fig. 5A).

**Figure 5.**
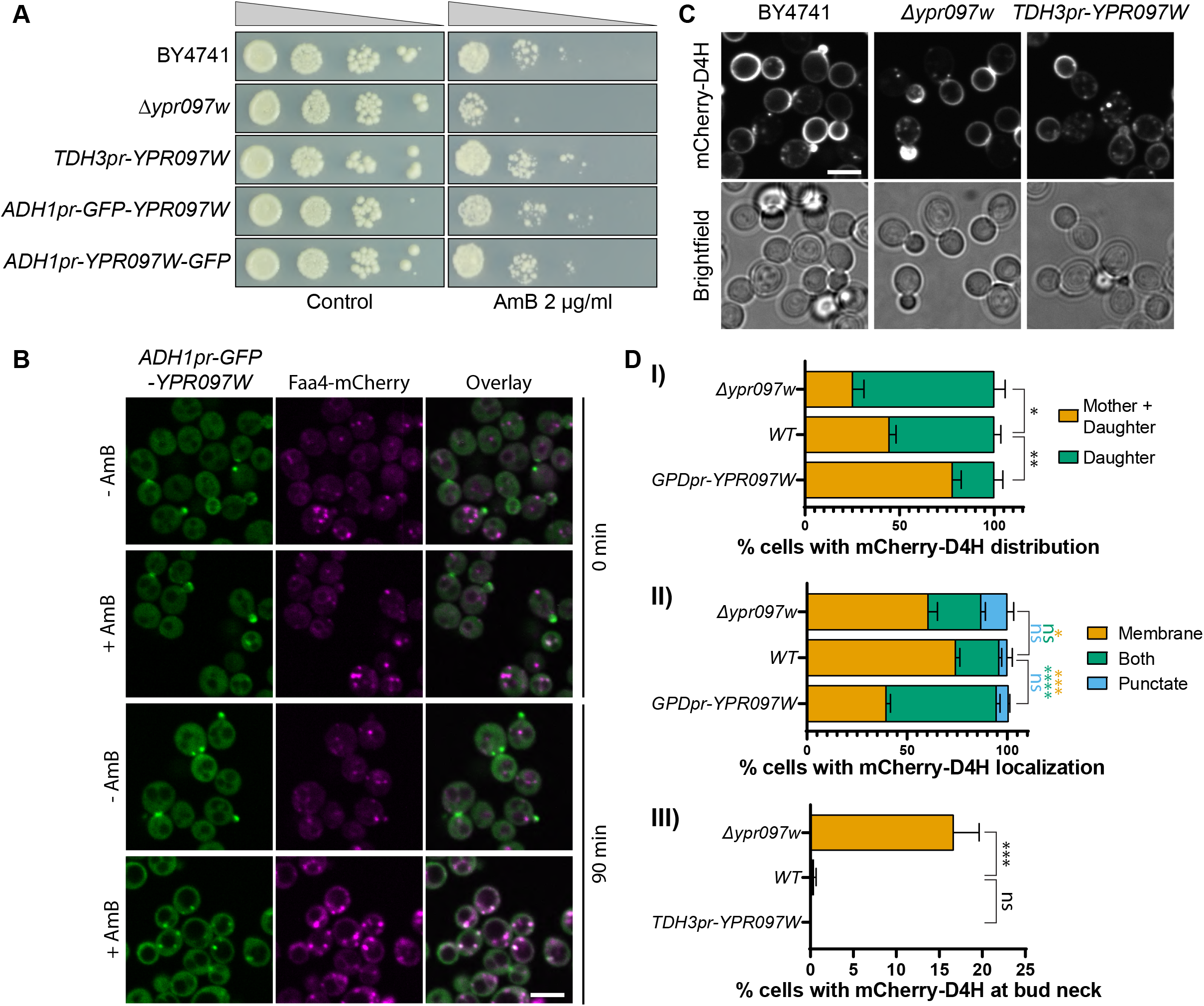
Ypr097w affects the cellular distribution of ergosterol. (**A**) Serial dilution growth assay in the presence/absence of Amphothericin B (AmB) (2 µg/ml), after 3 days of growth. (**B**) Localization of *ADH1pr-GFP-YPR097W* in control (DMSO) and AmB treated cells (2 µg/ml), at 0 min and after 90 min continuous drug exposure. Scale bar, 5 µm; Images obtained using Micro 2. (**C**) Cellular localization of the free ergosterol marker mCherry-D4H in WT, *Δypr097w* and overexpression *TDH3pr-YPR097W* cells. Cells were categorized based on marker distribution (mother+daughter cells or daughter) and localization (membrane, puncta or both), as well as accumulation at the bud neck. Scale bar, 5 µm; Images obtained using Micro 2. (**D**) Quantification of 3 independent experiments relative to (C). Data are presented as mean ± SEM (n=3). Analysis performed using ordinary one-way ANOVA with Dunnett’s multiple comparison test. ns. not significant, * p ≤0.05, ** p ≤0.01, *** p ≤0.001, **** p ≤0.0001.

Since we have previously seen that Ypr097w is highly dynamic in its localization we wondered if the cellular trigger for dynamics could be the free ergosterol levels on the PM. To assay this, we tracked the localization of GFP-Ypr097w in cells treated with AmB for 90 min and found that indeed, this resulted in a redistribution of Ypr097w (Fig. 5B). Under these conditions, Ypr097w almost exclusively localized to LDs (as marked by Faa4-mCherry). These results suggest that Ypr097w reacts to changes in PM composition and in turn, may affect the distribution of ergosterol in the cell, the levels of free ergosterol, or its capacity to be mobilized.

To track ergosterol distribution in cells, we constitutively expressed an optimized fragment of the bacterial toxin perfringolysin O, D4H, bound to a mCherry fluorophore. This sensor has been shown to bind free sterols on membranes with a concentration of at least 20 mol% sterols (Johnson et al., 2012; Maekawa and Fairn, 2015) and binds sterols at the PM in both fission and budding yeast (Kishimoto et al., 2021; Marek et al., 2020). In WT cells, this sterol reporter is mostly localized to the PM, with an enhanced signal at the daughter cell (Fig 5C). Additionally, small punctate structures can also be observed in some cells, generally in proximity to the PM (for examples of phenotypes, see Fig. 5 S1E). When *YPR097W* was either deleted or overexpressed, we observed striking effects on the distribution of the mCherry-D4H sterol reporter. In the absence of Ypr097w, a stronger localization to daughter cells was observed, with the additional accumulation of signal at the bud neck, a phenotype that is mostly absent from WT cells (Fig 5C, 5D(III)). This shift in localization is quite intriguing since Ypr097w itself is localized at the bud and bud neck in many cells. This could suggest that the presence of Ypr097w either shifts the distribution of ergosterol on the membrane, or protects it from interacting with mCherry-D4H. In cells where *YPR097W* is strongly overexpressed under the *TDH3* promoter, the distribution of mCherry-D4H was less polarized, with many cells displaying a similar signal between mother and daughter cells. Additionally, these cells showed a much stronger punctate or mixed pattern than both WT and *Δypr097w,* suggesting that overexpression of this protein affects the distribution of accessible ergosterol. Thus, while Ypr097w does not appear to regulate total ergosterol and sterol ester levels (Fig. 5 S1C-D), it appears to play a significant role in ergosterol localization in the cell.

Due to its distribution and effect on ergosterol we recommend the name Lec1 (<u>L</u>ipid-droplet <u>E</u>rgosterol <u>C</u>ortex 1) for Ypr097w.

## Discussion

For the past decade, there has been a growing interest in characterizing membrane contact sites and understanding their functions. This has been fueled by the development of new imaging and biochemical techniques and by the realization that contact site malfunction underlies multiple diseases (Castro et al., 2018; Herker et al., 2021; Scorrano et al., 2019; Zung and Schuldiner, 2020). However, identifying new contact sites and the proteins involved in their tethering and function still represents an important challenge in the field. Here, by taking advantage of split-Venus contact site reporters and high throughput techniques, we have significantly extended the pool of potential contact site resident proteins and effectors for six different contact sites in yeast. Of note is that our screens were far from being saturating since we only screened ∼1/3 of yeast proteins, albeit those with the highest chance of being direct residents and effectors. Even with only a subset of yeast proteins, our screens identified a large number of interesting candidates. Screening the additional proteins in the future may uncover more distal regulators such as enzymes that post-translationally modify contact site proteins or dedicated transcription factors. As a clear example of the effectiveness of this technique, we have detected previously known contact site proteins at the NVJ (Nvj1, Mdm1, Nvj3 and Lam6), Mdm1 at the LiDER, and Pex34 and Fzo1 at the PerMit. We anticipate that the newly reported residents and regulators will help drive the characterization of the three least studied contact sites – PerVale, pCLIP and GoPo.

We used this screening approach to identify potential roles for poorly characterized or uncharacterized proteins and report a new family of Vps13 related proteins: Hob1 (formerly Fmp27), Hob2 (formerly Ypr117w) and Csf1. Similar to Vps13, these proteins may localize to multiple contact sites. The localization of Hob1, Hob2 and Csf1 to both pCLIP and LiDER contacts underscores a possible function for this related family of proteins at LDs that may be shared by the human VPS13 and ATG2 homologs, which also localize to LDs (Kumar et al., 2018; Tamura et al., 2017; Velikkakath et al., 2012). Thus, these newly discovered members of the Vps13 family could potentially play similar roles in lipid transport between membranes. This function appears to be conserved in *Drosophila*, where a single homologue of the Hob1 and Hob2 proteins was initially described to play a role in intracellular trafficking (Neuman and Bashirullah, 2018). Furthermore, Csf1 was recently shown to function in lipid homeostasis under cold stress and in cells with rewired glycerophospholipid synthesis pathways (Peter et al., 2021). It is interesting to speculate that like Vps13, this set of related proteins may similarly function in glycerophospholipid transport at contact sites (Kumar et al., 2018; Li et al., 2020). Additionally, these proteins have homologs in mammalian cells – KIAA0100/BCOX1 for Hob1/2, and KIAA1109 for Csf1. Both of these have been implicated in human disease (Gueneau et al., 2018; Kane et al., 2019; Liu et al., 2014; Song et al., 2006).

While exploring the LiDER contact, we identified Ypr097w/Lec1, a previously uncharacterized protein. This protein stood out as it has a PX domain for binding lipid headgroups, as well as a FFAT motif (Loewen et al., 2003; Slee and Levine, 2019). Despite a clear dependence on yeast VAP proteins for its cellular distribution, we were unable to establish a role for the FFAT motif in directing Lec1 localization. However, we determined that Lec1 appears to act as a sensor of nutrient status, changing its cellular localization under multiple metabolic conditions, highlighting the ability of our screen to identify even transient contact site residents.

A recent *ab-initio* predicted structure of Ypr097w (AlphaFold2; (Jumper et al., 2021)) shows Lec1 is composed primarily of alpha helices that fold into a globular structure, with the PX domain on one side. The N- and C-terminal regions of the protein are highly interwoven which may explain why N- and C-terminal truncations disrupt its localization. Strikingly, the predicted structure of this protein shows a large internal hydrophobic cavity, with multiple channels opening to the cytoplasm. The hydrophobicity of the residues lining the pocket is highly conserved. This tantalizing structure suggests that Lec1 may represent a novel fold that binds or transports lipids.

The finding of an all helical protein forming a lipid binding cavity is not new. Such a cavity has been described for the insect allergen repeat domain, which contains two repeats of a domain of 5 helices, forming two hemispheres with one lipid inside (Mueller et al., 2013). The Lec1 cavity is uniquely large for a globular protein, at 29 nm^3^ almost 10-fold larger than any known cavity with a predominantly hydrophobic surface (Chwastyk et al., 2020). Since Lec1 is associated with membranes, lipid droplets and also with lipid modifying enzymes, we consider it possible that it encloses hydrophobic molecules such as lipids or intermediates of lipid synthesis. The cavity volume is over an order of magnitude bigger than the volume of a single lipid molecule. Hence we hypothesize that Lec1 could potentially accommodate a mixture of many different neutral lipids (e.g. triolein 1.62 nm^3^) and amphipathic lipids (e.g. DOPC 1.3 nm^3^). Interestingly, we found two additional proteins that share this predicted fold (using AlphaFold2) - Mdm1 and Nvj3, which fold together to form large spherical domains with an interior hydrophobic surface (Fig. 4E and G) that is highly conserved (Fig. 4F and H). While the PXA/nexin-C structure is similar to Lec1, they share no detectable sequence homology suggesting that other proteins with such a fold may exist. A more complete survey of predicted structures may reveal yet more proteins with large hydrophobic cavities that may function in lipid binding or transport.

We found that *Δlec1* cells are sensitive to AmB, an antifungal drug that interacts with free ergosterol at the PM and is suggested to induce membrane permeabilization and ergosterol removal (Anderson et al., 2014; Gray et al., 2012). Sensitivity to AmB has also been observed for other contact site proteins, such as Lam1, Lam2 (Ysp2) and Lam3 (Sip3), which localize to ER-PM contact sites and are involved in the retrograde transport of ergosterol from the PM to the ER (Gatta et al., 2015). By taking advantage of an ergosterol-binding fluorescent probe, we show that changes in the levels of Lec1 affect the localization of ergosterol in the cell, regulating its distribution between mother and daughter cells as well as internal reservoirs. The stronger accumulation of accessible ergosterol at the bud and bud neck of cells lacking Lec1 could explain why these mutants are more sensitive to AmB, as higher levels of accessible ergosterol would make budding cells more sensitive to this drug. Together with the lack of changes in overall ergosterol levels in these cells, our results suggest that Lec1 plays a role in the transbilayer organization or mobilization of ergosterol between membranes in yeast. Consistent with this, upon addition of AmB that binds accessible ergosterol, Lec1 shifts from buds and bud necks to LDs where it may either be sequestered or mobilize sterols or SE. Further investigation will be necessary to identify the mechanisms that regulate Lec1 localization in cells and to understand how this affects ergosterol distribution.

More generally our results show that despite the large volume of work on contact sites, multiple contact residents and regulators still await exploration. We anticipate that our work will provide a rich ground for further exploration of the contact site machinery.

## Materials and methods

### *S. cerevisiae* strains and plasmids

*S. cerevisiae* strains, plasmids and primers used in this study are described in Tables S5-7. Yeast strains were constructed from the laboratory strain BY4741 (Brachmann et al., 1998). Cells were genetically manipulated using the lithium acetate, polyethylene glycol (PEG) and single-stranded DNA (ssDNA) method for transformation (Gietz and Woods, 2006). Strains created using organelles markers expressed from plasmids were made fresh for imaging. Primers for genetic manipulations and validation were designed using Primers-4-Yeast (Yofe and Schuldiner, 2014). Plasmids expressing Ypr097w-GFPEnvy truncations were made by homologous recombination in yeast by co-transforming linearized plasmids with generated PCR products. The plasmids were recovered in *Escherichia coli* and sequenced. Plasmid MS1060 expressing GFP-Ypr097w WT was generated by isolation of the gene including 3’ and 5’ UTR by digestion of plasmid MD440 with ApaI and SacI, and cloning to plasmid MS889. Addition of N-terminal yeGFP was performed by restriction free cloning using primers 1060_ GFP_F and 1060_GFP_R, and plasmid MS246. Plasmids expressing Ypr097w phosphomutants were generated by overlap PCR and cloning using XbaI and NsiI restriction enzymes. In short, a fragment from plasmid MS1060 was amplified using the forward primer “YPR_seq2F” and three different reverse primers: 1065_S451E_NsiI_R, 1066_S451D_NsiI_R and 1067_S451A_NsiI_R, one for each mutation, respectively. PCR products and the original plasmid MS1060 were digested using XbaI and NsiI, and ligated to generate the three mutant plasmids. All oligonucleotides used in this study to generate PCR products for cloning are listed in Table S7.

### Growth conditions and microscopy

Yeast cells were cultured overnight in synthetic minimal media (SD) (0.67% [w/v] yeast nitrogen base with ammonium sulfate, 2% [w/v] glucose, amino acid supplements) at 30°C, unless stated otherwise. For microscopy experiments, cells were diluted and grown until mid-logarithmic phase. Cells were transferred to glass-bottom 384-well microscope plates (Matrical Bioscience) coated with concanavalin A (Sigma-Aldrich). After 20 min, wells were washed twice with media to remove non-adherent cells. Re-imaging of contact site screen hits was performed at room temperature (RT) using a VisiScope Confocal Cell Explorer system composed of a Zeiss Yokogawa spinning disk scanning unit (CSU-W1) coupled with an inverted Olympus IX83 microscope (named Micro1 for reference). Single focal plane images were acquired with a 60× oil lens and were captured using a PCO-Edge sCMOS camera, controlled by VisiView software (GFP/Venus at 488 nm, RFP/mCherry at 561 nm, or BFP/CFP at 405 nm). Additional imaging was performed using two additional microscopy systems (Micro 2 or 3; highlighted in figure legends). Micro 2 – an automated inverted fluorescence microscope system (Olympus) harboring a spinning disk high-resolution module (Yokogawa CSU-W1 SoRa confocal scanner with double micro lenses and 50 μm pinholes). Cells were recorded at 30°C using a 60X oil lens (NA 1.42) and with a Hamamatsu ORCA-Flash 4.0 camera. Fluorophores were excited by a laser and images were recorded in three channels: GFP (excitation wavelength 488 nm, emission filter 525/50 nm), mCherry (excitation wavelength 561 nm, emission filter 617/73 nm) and DAPI (excitation wavelength 405 nm, emission filter 447/60). Image acquisition was performed using scanR Olympus soft imaging solutions version 3.2. Images were transferred to ImageJ (https://imagej.nih.gov), for slight, linear, adjustments to contrast and brightness. Micro 3 - a DMi8 microscope (Leica Microsystems) with a high-contrast Plan Apochromat 63×/1.30 Glyc CORR CS objective (Leica Microsystems), an ORCA-Flash4.0 digital camera (Hamamatsu Photonics) and MetaMorph 7.8 software (MDS Analytical Technologies).

### Organelle labelling and drug treatment

To label LDs, cells were stained with BODIPY™ 493/503 (Invitrogen). For imaging after seeding in glass-bottom plates, cells were incubated with 1 µM of BODIPY (dissolved in DMSO) in SD media for 15min at RT and washed twice before imaging in SD media (control cells were treated with the same concentration of DMSO).

For amphotericin B (AmB, Invitrogen) treatment, cells were grown for 4h until mid-log, seeded on glass-bottom plates and imaged once before adding AmB. AmB was added to SD media at 2 µg/ml and cells were incubated and imaged every 30min at 30°C.

For nutrient shift experiments, cells were first imaged in minimal synthetic dextrose media, the media aspirated and replaced with PBS with or without glucose supplementation.

### High content contact site protein screens

To identify new potential contact site proteins, query strains containing contact site reporters for six different contact sites were crossed against a subset of the SWAT *TEF2-mCherry* library (1165 strains) (Weill et al., 2018; Yofe et al., 2016) using the synthetic genetic array method (Cohen and Schuldiner, 2011; Tong and Boone, 2006). This subset of the SWAT library contains all strains where protein localization is annotated as “punctate”, as this is the most common phenotype for contact site proteins, as well as a set of known contact site proteins. Protein punctate localization was based on previous annotations manually performed in our laboratory for three different libraries: *NOP1*-GFP, *TEF2*-mCherry and NATIVEpr-GFP, and the equivalent *TEF2*-mCherry strains annotated as punctate in both NOP1-GFP and NATIVEpr-GFP were also added to the final subset. To perform yeast manipulations, query and library strains were handled in high-density format (384-1536 strains per plate) using a RoToR bench-top colony arraying instrument (Singer Instruments, UK). In short, cells were mated on rich medium plates and diploids were selected in SD_MSG_-His containing Geneticin (200 µg/ml) (Formedium) and Nourseothricin (200 µg/ml) (WERNER BioAgents “ClonNat”). Sporulation was induced by transferring cells to nitrogen starvation media plates for 8 days. Haploid cells were selected in SD-Leu (for Mat alpha selection) and -Arg-Lys with toxic amino-acid derivatives canavanine and thialysine (Sigma-Aldrich) to select against diploids. Finally, haploid cells containing the combination of manipulations desired were selected using SD_MSG_-His containing Geneticin (200 µg/ml) and Nourseothricin (200 µg/ml), and final libraries containing the genomic traits were created. For each library, a set of strains were verified by microscopy and check PCR.

The new libraries were screened using an automated microscopy setup. Cells were transferred from agar plates into 384-well plates for growth in liquid media using the RoToR arrayer. Liquid cultures were grown in a LiCONiC incubator overnight at 30 °C in SD-His. A JANUS liquid handler (PerkinElmer) connected to the incubator was used to dilute the strains to an OD600 of ∼0.2 into plates containing SD-His medium, and plates were incubated at 30 °C for 4 h. Strains were then transferred by the liquid handler into glass-bottom 384-well microscope plates (Matrical Bioscience) coated with Concanavalin A (Sigma-Aldrich). After 20 min, wells were washed twice with SD-Riboflavin media to remove non-adherent cells and to obtain a cell monolayer. The plates were then transferred to an Olympus automated inverted fluorescent microscope system using a robotic swap arm (Hamilton). Cells were imaged in SD-Riboflavin at 18-20 °C using a ×60 air lens (NA 0.9) and with an ORCA-ER charge-coupled device camera (Hamamatsu), using the ScanR software. Images were acquired in two channels: GFP (excitation filter 490/20 nm, emission filter 535/50 nm) and mCherry (excitation filter 572/35 nm, emission filter 632/60 nm). After acquisition, images were manually reviewed using ImageJ. Hits form the library were re-picked and re-imaged using a spinning disk confocal microscope (as above, Micro 1) to improve resolution, and only proteins that still fit the co-localization/effect criteria were kept in the final hit list.

The final list of hits for each contact site was manually annotated. As some organelles are smaller than the diffraction limit of confocal microscopy (peroxisomes, lipid droplets and Golgi), we were unable to distinguish proteins localized to the organelle from those localized to the contact sites analyzed. To simplify the hit list, we assembled a list of known proteins for these organelles, and removed them if they were marked as co-localized in the final hit list unless they affected the contact site extent or brightness.

### Protein modelling

Ab initio folding by co-evolution analysis by trRosetta and Alphafold2 was carried out at the Yanglab web server and at the AlphaFold2.ipynb colab respectively (Mirdita et al., 2021; Yang et al., 2020). Settings were standard, with AlphaFold2 set to return relaxed models (Amber = On) and to ignore templates. For Ypr097w, uninformative regions were omitted in trRosetta 212-244 and 366-465; in AlphaFold2 305-321, 356-468 and 690-780. In the latter case, to model the complete protein, omitted regions were re-added using the new AlphaFold2 structure as the template, by one-to-one threading in Phyre2 (Kelley et al., 2015). To model Csf1, since neither it nor any homolog has been published among the proteins modelled by AlphaFold2, 4 overlapping sections (starting with residues 1-1080 and ending with 2070-2959) were predicted by the AlphaFold2.ipynb colab. A single model was created from overlapping portions in Chimera (Pettersen et al., 2004). Proteins were visualized in CCP4MG, Chimera (Pettersen et al., 2004)and ChimeraX (Pettersen et al., 2021) software. Cavity sizes were calculate with MOLE (Pravda et al., 2018).

### Phenotype quantification

For quantification of Ypr097w localization, 200 cells (mix of single and budding) were manually analyzed from each of 3 biological replicates (total n=3, 600 cells). For LD quantification, 100 cells (mix of single and budding) from 3 independent experiments were manually counted, from a single focal plane (total n=3, 300 cells). For mCherry-D4H localization and distribution, 100 cells (budding) from 3 independent experiments were analyzed (total n=3, 300 cells). To avoid quantification bias, cells were selected in the brightfield channel and then analyzed for GFP or mCherry localization. For comparison of LD number between WT, *Δypr097w* and *TDH3pr-YPR097W*, mCherry-D4H distribution, localization, and bud neck localization, an ordinary one-way ANOVA test was used with Dunnett’s correction for multiple comparisons. ns. not significant, * p ≤0.05, ** p ≤0.01, *** p ≤0.001, **** p ≤0.0001. Graphs and data analysis was performed using GraphPad Prism 9 (GraphPad Software Inc., San Diego, CA).

### Serial dilution growth assay

To assess sensitivity to AmB, cells were grown overnight in SD media and back diluted for 4h until they reached mid-logarithmic growth. For each strain, cells were diluted to a starting OD600=0.1 and serial 10-fold dilutions were prepared in 96-well plates. 2.5 µl of each dilution were plated in SD agar plates containing 2 µg/ml of AmB and imaged 3 days after growing at 30 °C. Plates without AmB were prepared in parallel to control for the effect of each genetic manipulation on growth. For each condition, 2 replicate plates were prepared in each of 3 independent experiments.

### Lipid droplet purification

Lipid droplets were purified as previously described (Athenstaedt, 2010) with modifications (Ganesan et al., 2020). Briefly, yeast cultures were grown for 24h in defined medium. Cells were then collected, washed once with water and the wet weight was determined. Cells were shaken at 30°C for 10 min in 0.1 M Tris/ H_2_SO_4_ buffer (pH 9.4) containing freshly added 10 mM DTT and then washed with 1.2 M sorbitol in 20 mM KH_2_PO_4_ (pH 7.4). Zymolyase 20T (2 mg per gram cell wet weight) was added to obtain spheroplasts (Daum et al., 1982). Spheroplasts were washed twice with 1.2 M sorbitol in 20 mM KH_2_PO_4_ (pH 7.4) prior to homogenization. The spheroplasts were then resuspended in buffer A (10 mM MES-Tris (pH 6.9) 12% (w/w) Ficoll 400–0.2 mM EDTA, 1 mM PMSF 1.5 μg/ml pepstatin and Complete EDTA-free protease inhibitor mixture) to a final concentration of 2.5 g per cell wet weight per ml. Spheroplasts were then homogenized using a Dounce homogenizer by employing 30 strokes using a loose-fitting pistil, followed by a centrifugation at 5,000 g at 4°C for 5 min. The resulting supernatant (∼5 ml) was transferred into an ultracentrifugation tube, overlaid with an equal volume of buffer A, and centrifuged at 100,000 g at 4°C for 60 min in a swinging rotor (SW40Ti) (Beckman Coulter OptimaTM L–100K). Following centrifugation, a white floating layer was collected from the top of the overlay and resuspended gently in buffer A by using a homogenizer with a loose-fitting pistil. The homogenized floating layer was again transferred into an ultracentrifugation tube, overlaid with buffer B (10 mM MES-Tris (pH 6.9) 8% (w/w) Ficoll 400–0.2 mM EDTA) and centrifuged at 100,000 g (4°C) for 30 min. After the second centrifugation, the top white floating layer containing LDs was removed. A subphase laying beneath the white floating layer was also collected (∼0.3 ml). The top white floating layer was then suspended in buffer containing 8% (w/w) Ficoll 400/0.6 M sorbitol and overlaid with buffer containing 0.25 M sorbitol. After a centrifugation at 100,000 g for 30 minutes, a final white floating layer was collected containing highly purified LDs. The purified LDs were homogenized using a Dounce homogenizer by applying 20 strokes.

### Western blot analysis of lipid droplet fractions

BCA assay (Thermo Scientific) with bovine serum albumin as a standard was used to determine protein concentration. Samples were resuspended in gel loading buffer (0.2 M Tris HCl (pH 6.8), 8% SDS, 0.4% bromophenol blue, and 40% glycerol) and boiled for 1 minute. SDS-PAGE was carried out by the Laemmli method (Laemmli, 1970). Proteins were separated by 8% resolving gel containing trichloroethanol (TCE, Sigma) to visualize proteins (Ladner et al., 2004). Proteins were transferred to a polyvinylidene fluoride (PVDF) membrane (Millipore) using a Bio-Rad transfer system at 90 V for 80 min, and then stained with Red Ponceau (Sigma) to monitor the transfer. For western blot analysis, monoclonal anti-GFP antibody (Roche) and polyclonal antibody raised against Erg6 (kind gift from G. Daum, Universität Graz) were used and subsequently with horseradish peroxidase-conjugated secondary antibodies (Invitrogen, Thermo Fisher). Enhanced chemiluminescence (Amersham, GE Healthcare) was used for detection followed by imaging (Amersham Imager 600).

### Lipid extraction and thin layer chromatography

The indicated yeast cultures were grown for 24h in synthetic defined medium. Cultures were then diluted and allowed to grow into mid-log phase (5 h), early stationary phase for an additional 24 hours or additional 48 h for late stationary phase. For growth resumption samples, cells were pelleted after growing for 24 h, resuspended in fresh medium and cultured for an additional 45 min. For each of the growth phases, 20 OD600 of cells were collected. Lipid extractions were performed as previously described (Zaremberg and McMaster, 2002). Briefly, cell pellets were resuspended in 1 ml of CHCl_3_/CH_3_OH (1/1), and disrupted with 0.5 mm acid-washed glass beads for 1 min at 4 °C using a BioSpec Multi-Bead Beater. The supernatant was transferred into a glass tube and the beads were washed with 1.5 ml of CHCl_3_/CH_3_OH (2/1, v/v) and transferred into the same glass tube. A total of 1.5 ml of water and 0.5 ml of CHCl_3_ was added to the glass tube to facilitate phase separation. The organic phase containing the lipids was collected after a 10 min centrifugation at 2500 g. A total of 2.5 ml of artificial aqueous layer, CHCl_3_/CH_3_OH/H_2_0 (3/48/47, v/v/v) was then added to the organic phase and subjected to another 10 min centrifugation at 2500 g. The final organic phase was collected and dried using nitrogen gas. Neutral lipids were analyzed by thin layer chromatography using Silica Gel on TLC aluminum foil plates (Sigma) and separated using solvent system I: petroleum ether/diethyl ether/acetic acid (80/20/1, v/v/v). Lipids were resuspended in CHCl_3_ and equal amount of lipids were spotted on the plates. Lipids were visualized using iodine vapor staining or by dipping the plates in a solution containing 3.2% H_2_SO_4_, 0.5% MnCI_2_, 50% EtOH and charring the plates at 120°C for 30 min. Plates were imaged using an Amersham Imager 600. Neutral lipid quantification was done using the linear range of lipid standards and ImageJ. GraphPad Prism 5 software was used for statistical analysis of data and preparation of figures.

Lipid standards used were purchased from Sigma: cholesteryl-ester (700222), 1-2-dioleoyl-sn-glycerol (800811), ergosterol (E6510), oleic acid (O1008), squalene (S3626), and triolein (870110).

### Cell culture and harvest protocol for lipidomics

A single colony from an agar plate with synthetic complete dextrose (SCD) medium was used to inoculate a pre-culture of 3 mL in liquid SCD medium. After cultivation under constant agitation at 30°C for 19 h, stationary cells were used to inoculate a main culture to an OD600 of 0.1. Logarithmically growing cells were harvested at an OD600 of 1.0, while stationary cells were harvested 48 h after inoculation of the main culture. 20 OD units of cells were harvested by centrifugation (3,500 g, 5 min, 4°C), followed by three washing steps in ice-cold buffer containing 155 mM ammonium bicarbonate and 10 mM sodium azide using rapid centrifugation (10,000 g, 20 s, 4 °C). The final cell pellet was snap-frozen with liquid nitrogen and stored at -80°C until cell lysis. To this end, the cell pellets were thawed on ice and then resuspended in 1 mL of ice-cold 155 mM ammonium bicarbonate. The suspension was transferred into a fresh 1,5 mL reaction tube containing 200 µL of zirconia beads (0,5 mm bead size). Cell disruption was induced by vigorous shaking using a DisruptorGenie for 10 min at 4°C. 500 µL of the resulting lysate was snap-frozen and sent to Lipotype GmbH for lipid extraction and further analysis.

### Lipid extraction for mass spectrometry (lipidomics)

Mass spectrometry-based lipid analysis was performed by Lipotype GmbH (Dresden, Germany) as described (Ejsing et al., 2009; Klose et al., 2012). Lipids were extracted using a two-step chloroform/methanol procedure (Ejsing et al., 2009). Samples were spiked with internal lipid standard mixture containing: CDP-DAG 17:0/18:1, cardiolipin 14:0/14:0/14:0/14:0 (CL), ceramide 18:1;2/17:0 (Cer), diacylglycerol 17:0/17:0 (DAG), lyso-phosphatidate 17:0 (LPA), lyso- phosphatidyl- choline 12:0 (LPC), lyso-phosphatidylethanolamine 17:1 (LPE), lyso- phosphatidylinositol 17:1 (LPI), lyso-phosphatidylserine 17:1 (LPS), phosphatidate 17:0/14:1 (PA), phosphatidylcholine 17:0/14:1 (PC), phosphatidylethanolamine 17:0/14:1 (PE), phosphatidylglycerol 17:0/14:1 (PG), phosphatidylinositol 17:0/14:1 (PI), phosphatidylserine 17:0/14:1 (PS), ergosterol ester 13:0 (EE), triacylglycerol 17:0/17:0/17:0 (TAG), stigmastatrienol, inositolphosphorylceramide 44:0;2 (IPC), mannosyl-inositolphosphorylceramide 44:0;2 (MIPC) and mannosyl-di- (inositolphosphoryl)ceramide 44:0;2 (M(IP)2C). After extraction, the organic phase was transferred to an infusion plate and dried in a speed vacuum concentrator. 1st step dry extract was re-suspended in 7.5 mM ammonium acetate in chloroform/methanol/propanol (1:2:4, V:V:V) and 2nd step dry extract in 33% ethanol solution of methylamine in chloroform/methanol (0.003:5:1; V:V:V). All liquid handling steps were performed using Hamilton Robotics STARlet robotic platform with the Anti Droplet Control feature for organic solvents pipetting.

### Mass spectrometry data acquisition (lipidomics)

Samples were analyzed by direct infusion on a QExactive mass spectrometer (Thermo Scientific) equipped with a TriVersa NanoMate ion source (Advion Biosciences). Samples were analyzed in both positive and negative ion modes with a resolution of Rm/z=200=280000 for MS and Rm/z=200=17500 for MSMS experiments, in a single acquisition. MSMS was triggered by an inclusion list encompassing corresponding MS mass ranges scanned in 1 Da increments (Surma et al., 2015). Both MS and MSMS data were combined to monitor EE, DAG and TAG ions as ammonium adducts; PC as an acetate adduct; and CL, PA, PE, PG, PI and PS as deprotonated anions. MS only was used to monitor LPA, LPE, LPI, LPS, IPC, MIPC, M(IP)2C as deprotonated anions; Cer and LPC as acetate adducts and ergosterol as protonated ion of an acetylated derivative (Liebisch et al., 2006).

### Data analysis and post-processing (lipidomics)

Data were analyzed with in-house developed lipid identification software based on LipidXplorer (Herzog et al., 2012, 2011). Data post-processing and normalization were performed using an in-house developed data management system. Only lipid identifications with a signal-to-noise ratio >5, and a signal intensity 5-fold higher than in corresponding blank samples were considered for further data analysis.

For statistical analysis of lipidomics data, the mol % of each lipid class was calculated against the total sum of lipids per sample. For each metabolic state (mid-logarithmic growth and stationary), 3 biological replicates of WT vs *THD3pr-YPR097W* strains were compared using multiple unpaired t-tests, assuming Gaussian distribution. Data was corrected for False Discovery Rate using the two-stage step-up method of Benjamini, Krieger and Yekutieli (Q=5%). Data analysis was performed using GraphPad Prism 9 (GraphPad Software Inc., San Diego, CA).

### DHFR Protein Fragment Complementation Assay and Ontology Analysis

Endogenous YPR097W was C-terminally fused to one half of a methotrexate-resistant variant of the essential DHFR enzyme in a MATa strain, which was crossed into a library of MATα strains (n = ∼4300) expressing proteins fused to the complementary half of the DHFR enzyme (Tarassov et al., 2008). Diploids were subjected to two rounds of double mutant selection followed by two rounds of selection on media containing 200 µg/µl methotrexate (MTX). All yeast manipulations were carried out using the BM3-BC microbial pinning robot (S&P Robotics Inc., Canada) with a 384-pin tool. Colonies were maintained in either 768- or 1536-density and technical duplicates were pinned for MTX selection steps. Colony area was analyzed using CellProfiler (Lamprecht et al., 2007), after 7 days at 30 °C. Interactions with abundant, cytosolic proteins were filtered out by comparing colony area of Ypr097w DHFR diploids to DHFR diploids containing an overproduced DHFR bait fragment fused to a cytosolic reporter. Z-scores were generated using median colony area from two technical replicate for each Ypr097w-prey combination. Functional analysis of Ypr097w DHFR interactors (Z>2.5) was performed using the Gene Ontology (Ashburner et al., 2000; The Gene Ontology Consortium, 2019) GO Enrichment Analysis tool (Mi et al., 2019).

## Acknowledgments and Funding

We wish to thank Einat Zalckvar and Maria Bohnert for enriching scientific discussions, many great ideas and critical reading of this manuscript. We thank Yoav Peleg for cloning of the phosphomutant plasmids. We thank Ron Rotkopf for statistical analysis support. We thank Christian Landry for providing the C-terminally tagged DHFR prey library and plasmid for generation of a bait Ypr097w construct. We thank Joel Goodman, Sophie G. Martin, Won-Ki Huh, David Breslow, Naama Barkai and Vlad Denic for plasmids.

Collaborative work between the Ernst and Schuldiner labs is supported by a VolksWagen (VW) Foundation “Life” grant (93092). Work in the Schuldiner lab is also supported by a Deutsche Forschungsgemeinschaft (DFG) SFB1190 grant. MS is an incumbent of the Dr. Gilbert Omenn and Martha Darling Professorial Chair in Molecular Genetics. IGC is a recipient of an EMBO Long-term Fellowship (ALTF-580-2017). CJS is supported by MRC funding to the MRC LMCB University Unit at UCL, award code MC_UU_00012/6. TL was funded by the BBSRC UK (Grant BB/M011801/1). The Conibear lab acknowledges support by the Canada Foundation for Innovation (Leading Edge Fund 30636); Canadian Institutes of Health Research (grant 148756 to EC, CGS-M Frederick Banting and Charles Best Canada Graduate Scholarship to SKD and SPS, CGS-D Frederick Banting and Charles Best Canada Graduate Scholarship to SKD); Natural Sciences and Engineering Research Council of Canada (PGS-D to SPS); BC Children’s Hospital Research Institute (Sue Carruthers Graduate Studentship to SKD and Jan M. Friedman Graduate Studentship to SPS) and University of British Columbia 4-Year Doctoral Fellowship to SKD and SPS. VZ is supported by the Natural Sciences and Engineering Research Council of Canada (NSERC).

## Table Legends

**Table S1. List of strains in the “Puncta Library” collection.** List of the subset of strains from the SWAT *TEF2pr*-mCherry library selected for this work based on protein localization in cells.

**Table S2. Newly suggested contact site residents and regulators.** List of hits for each of the 6 screens performed and their phenotype based on co-localization with contact site reporter and/ effect over the contact site reporter.

**Table S3. Ypr097w DHFR Proximity Interactome.** List of noise-filtered Ypr097w DHFR interactors (Z-Score>2.5). Hits with reported lipid droplet localization (data from (Huh et al., 2003; Weill et al., 2018) or membership in a validated LD proteome (data from (Currie et al., 2014)) are colored pink, while hits with reported ER localization that are not present in the validated LD proteome are colored blue.

**Table S4. Lipidomic analysis of WT and TDH3pr-YPR097W strains.**

**Table S5. *S. cerevisiae* strains used in this study.**

**Table S6. Plasmids used in this study.**

**Table S7. Primers used in this study.**

## Source data

**Zip folder – CastroEtAl_Source data_blots**

**Figure 3F anti-GFP-YPR097W_blot - source data 1 –** Original western blot from figure 3F, top panel.

**Figure 3F anti-GFP-YPR097W_blot - source data 2 -** Annotated western blot from figure 3F, top panel.

**Figure 3F anti-Erg6_blot - source data 1 -** Original western blot from figure 3F, bottom panel.

**Figure 3F anti-Erg6_blot - source data 2 -** Annotated western blot from figure 3F, bottom panel.

**Figure 5 - figure supplement 1C - Mid-log_Stationary 24h - source data 1 –** Thin layer chromatography, original blot from figure 5 – figure supplement 1C, left panel.

**Figure 5 - figure supplement 1C - Mid-log_Stationary 24h - source data 2–** Thin layer chromatography, annotated blot from figure 5 – figure supplement 1C, left panel.

**Figure 5 - figure supplement 1C - GR_Stationary 48h - source data 1–** Thin layer chromatography, original blot from figure 5 – figure supplement 1C, right panel.

**Figure 5 - figure supplement 1C - GR_Stationary 48h - source data 2–** Thin layer chromatography, annotated blot from figure 5 – figure supplement 1C, right panel.

## Excel files

**Figure 1B-source data –** data used for graph present in Figure 1B

**Figure 3C-source data –** data used for graph present in Figure 3C

**Figure 3D-E-source data –** data used for graph present in Figure 3D-E

**Figure 5D-source data –** data used for graph present in Figure 5D and statistical results

**Figure 5 - figure supplement 1A-source data –** data used for graph present in Figure 5 – figure supplement 1A and statistical results

**Figure 5 - figure supplement 1D-source data –** data used for graph present in Figure 5 – figure supplement 1D and statistical results

---

**Figure 1 S1.**
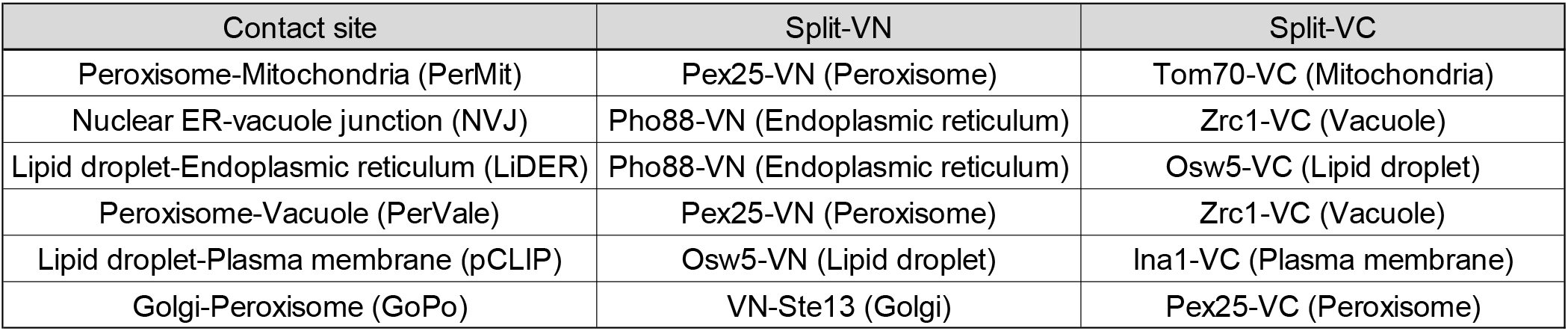
Proteins used for creation of the various split-Venus reporters. Table showing the pairs of tagged proteins used as reporters in each contact site screen, the Venus protein fragment with which they were fused, and the organelle in which they reside.

**Figure 2 S1.**
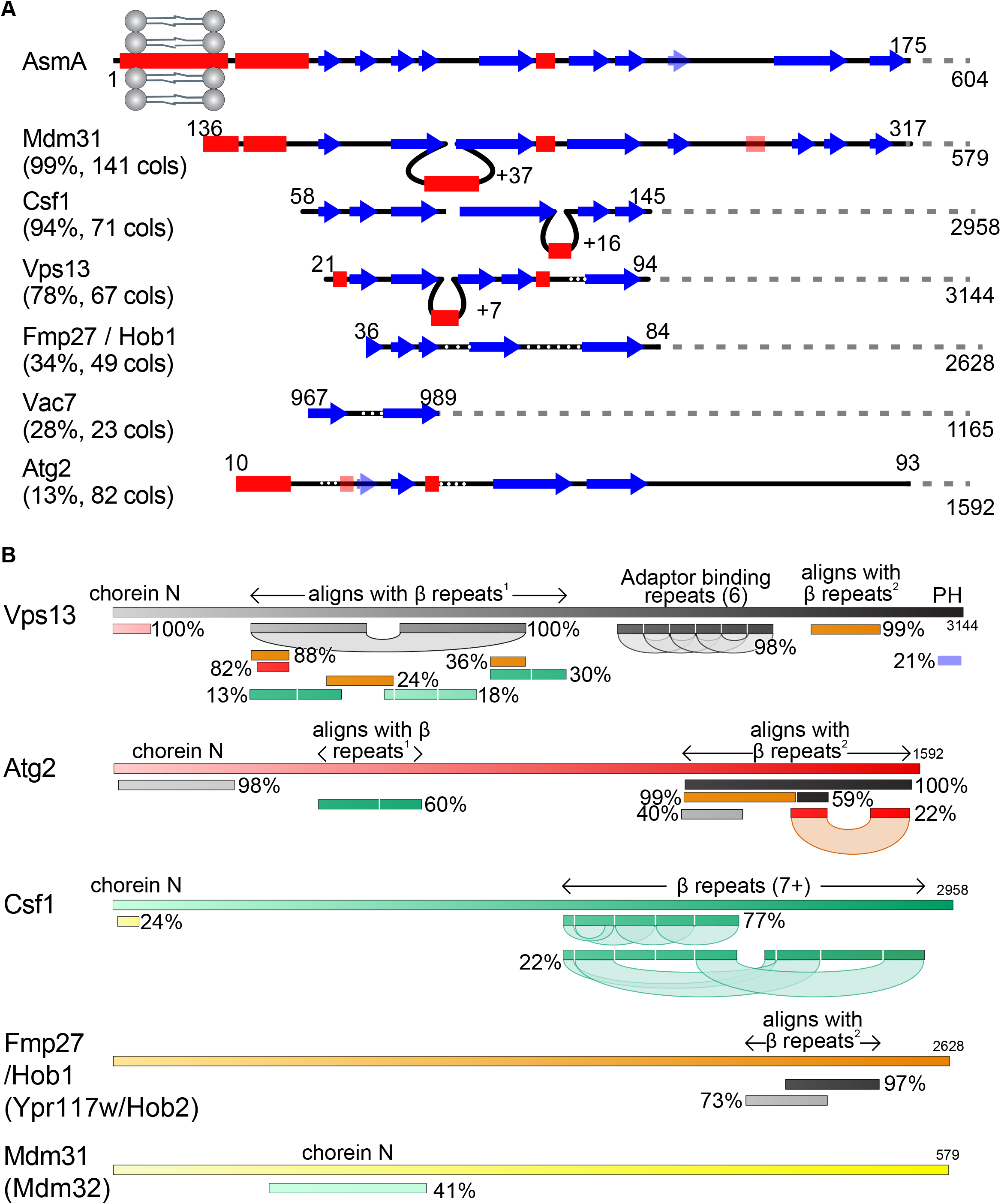
Vps13 family protein domain structure and comparison. (A) Identification of Csf1 and Hob1 as homologs of Vps13. The N-terminus of AsmA (*E. coli*), consisting of its transmembrane domain and N-chorein domain was used to seed an HHpred search for yeast homologs. Secondary structural elements are shown of the seed and all yeast hits that focus on the predicted β-strands with >15 columns matched. For each hit, the probability of the match being a true positive is shown together with the number of columns (cols) matched. Insertions in hits are indicated by loops, and gaps in hits are indicated by white dotted lines. Weakly predicted secondary structural elements are shown as faint and transparent. (B) Structural elements determined by HHpred in the superfamily of Vps13, Atg2, their newly described homologues Csf1, Hob1/2 (Fmp27/Ypr117w) and Mdm31/32. HHpred searches with each full-length protein produced alignments between proteins in the superfamily. In addition, self- alignments occurred in Vps13, Atg2 and Csf1, identifying three types of repeats, only one of which, the 6 Adaptor Binding repeats in the C-terminal half of Vps13, has been identified previously (Bean et al., 2018; Kumar et al., 2018). The most common type of repeat is a mainly-beta strand module of approx. 140 residues, which is repeated at least 7 times in the C-terminal half of Csf1. This region is homologous directly to regions in the N-terminal halves of Vps13 and Atg2, which in turn are homologous to regions near the C-termini of Vps13, Atg2 and Hob1/2. The final type of repeats consist of a pair of helices, which is repeated near the C-terminus of both Vps13 and Atg2, and has previously been named “Atg2_C”. Other domains (both identified previously) identified by homologies are: Chorein-N at the N-terminus of Vps13, Atg2, Csf1 and Mdm31/32, and the pleckstrin homology (PH) domain at the C-terminus of Vps13 only. Position along each protein is indicated in both seeds and hits by varying colors across different spectra. Hob2 (Ypr117w) and Mdm32 were merged with their paralogs as on this schematic level they are identical to their paralogs Hob1 (Fmp27) and Mdm31, respectively.

**Figure 2 S2.**
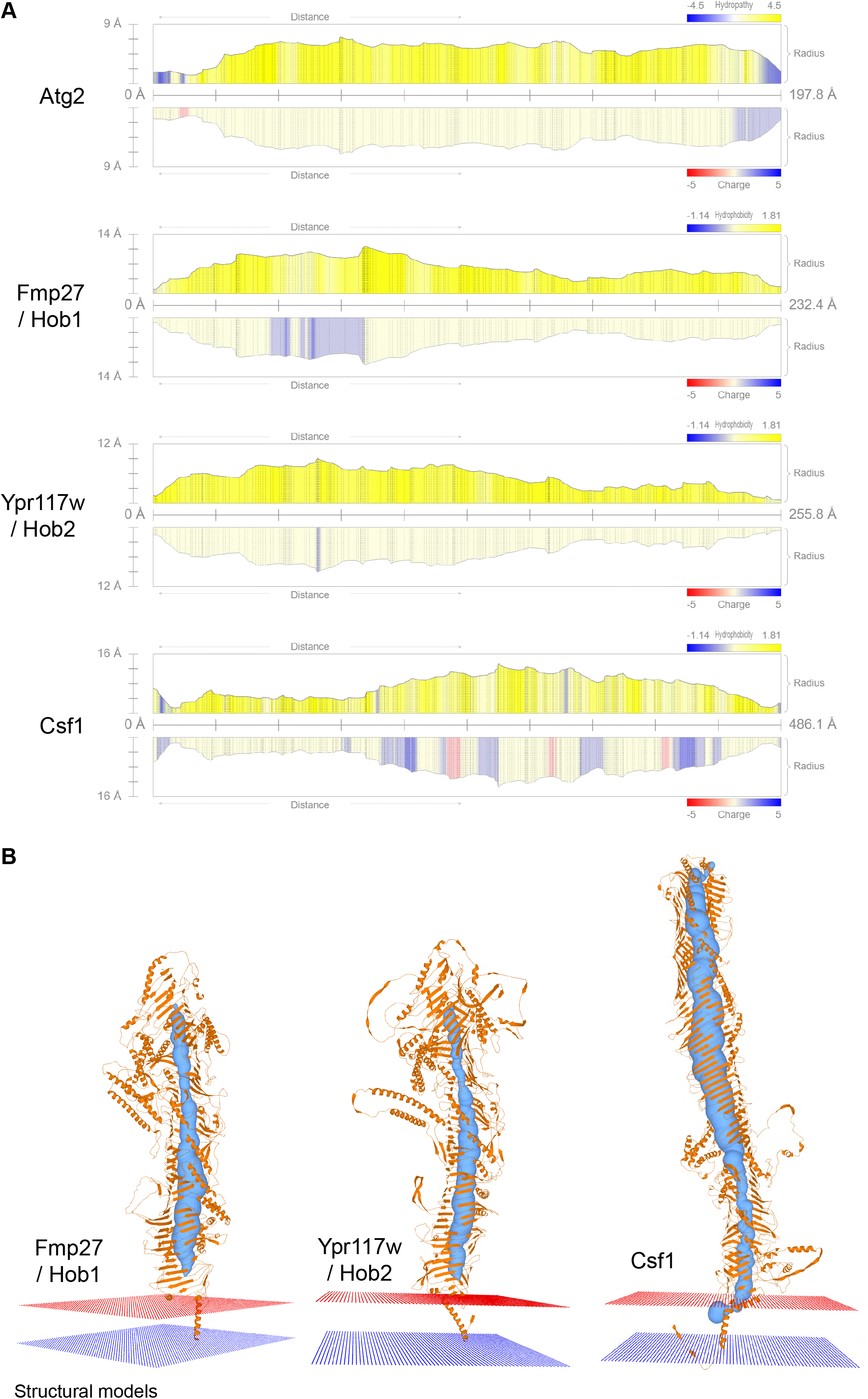
Vps13 family protein channel comparison and membrane topology. (A) Channel profiles obtained with MOLE for the different Vps13 family proteins. The radius of the channel along the protein is depicted, together with its hydrophobicity and charge. (B) Predicted N-terminal transmembrane domains anchors for the Vsp13 family proteins analyzed with MOLE.

**Figure 3 S1.**
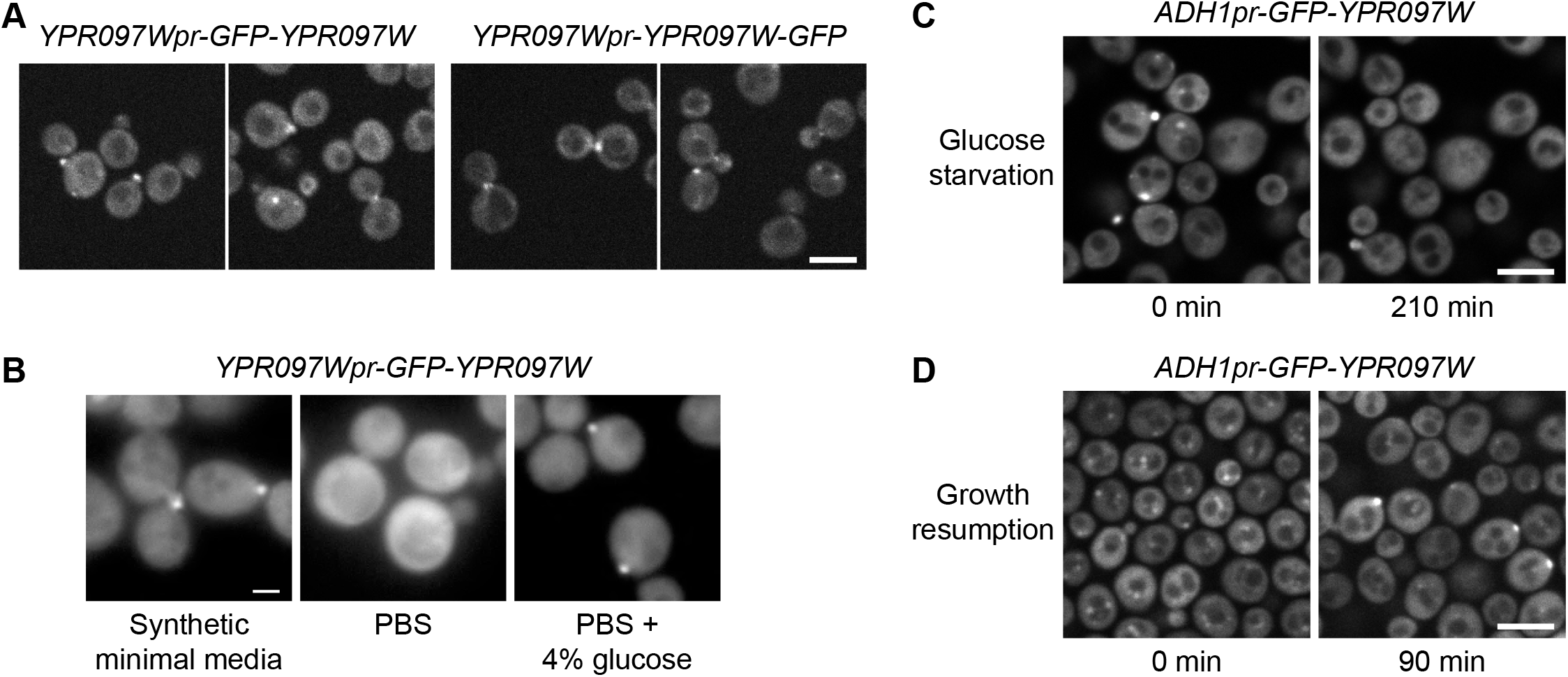
Ypr097w dynamics. (A) Localization of endogenous Ypr097w, GFP-tagged at the N-terminus (2 left images) or C- terminus (2 right images). Scale bar, 5 µm; Images obtained using Micro 1. (B) Loss of bud and bud neck localization of endogenous GFP-Ypr097w when shifted from minimal synthetic media to PBS (<2min). Bud and bud neck localization remains when PBS is supplemented with glucose. Scale bar, 2 µm; Images obtained using Micro 3. (C) Loss of bud/bud neck and puncta localization of *ADH1pr-GFP-YPR097W* when shifted to synthetic media without glucose for 210 min. Scale bar, 5 µm; Images obtained using Micro 2. (D) Cellular localization of *ADH1pr-GFP-YPR097W* in stationary (left) and upon media replenishment for 90 min (growth resumption, right). GFP-Ypr097w re-localizes from a punctate pattern to a combination of punctate, bud/bud neck and cytosol. Scale bar, 5 µm; Images obtained using Micro 2.

**Figure 3 S2.**
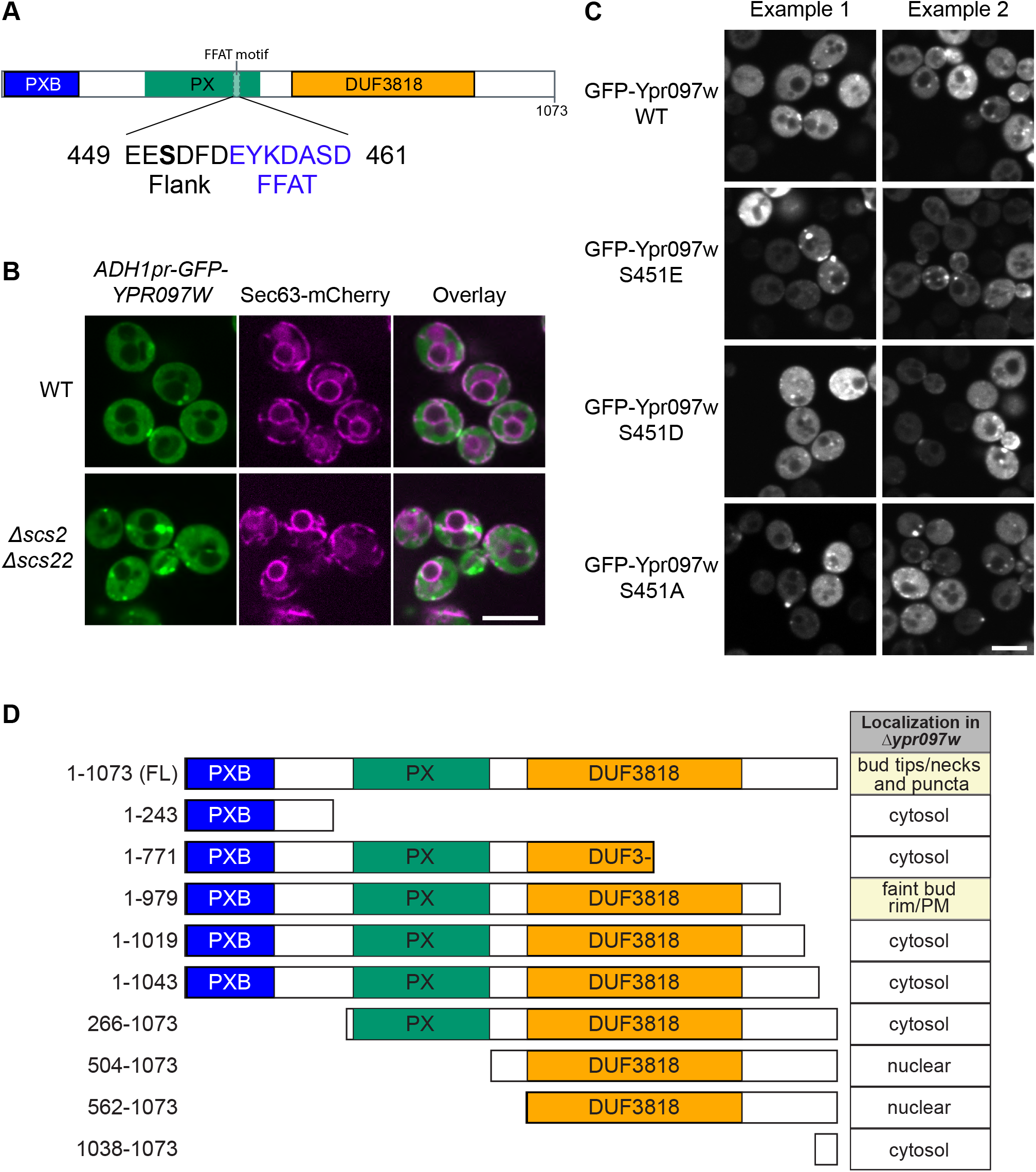
Effect of Ypr097w truncation or mutation on cellular localization. (A) Schematic representation of Ypr097w domains, with highlighted FFAT motif and previously detected phosphorylation site. 3 phosphomutants were generated by site directed mutagenesis of residue S451: phosphomimetic (S451E; S451D) and phospho-null (S451A). (B) Cellular localization of *ADH1pr-GFP-YPR097W* in control and *Δscs2Δscs22* cells. GFP- Ypr097w localization shifts to a stronger punctate pattern in *Δscs2Δscs22* cells. ER marker is Sec63-mCherry. Scale bar, 5 µm; Images obtained using Micro 2. (C) Cellular localization of GFP-Ypr097w WT and phosphomutants in *Δypr097w* cells. Plasmid expressed GFP-Ypr097w localizes to bud/bud neck and punctate pattern with different levels of expression. Two examples of each plasmid are shown to exemplify phenotype variability. Scale bar, 5 µm; Images obtained using Micro 2. (D) Summary of the cellular localization and schematic representation of the Ypr097w truncations examined by microscopy. Table indicates the localization of plasmid-borne C-terminally GFPenvy- tagged truncations in a Δ*ypr097w* strain.

**Figure 4 S1.**
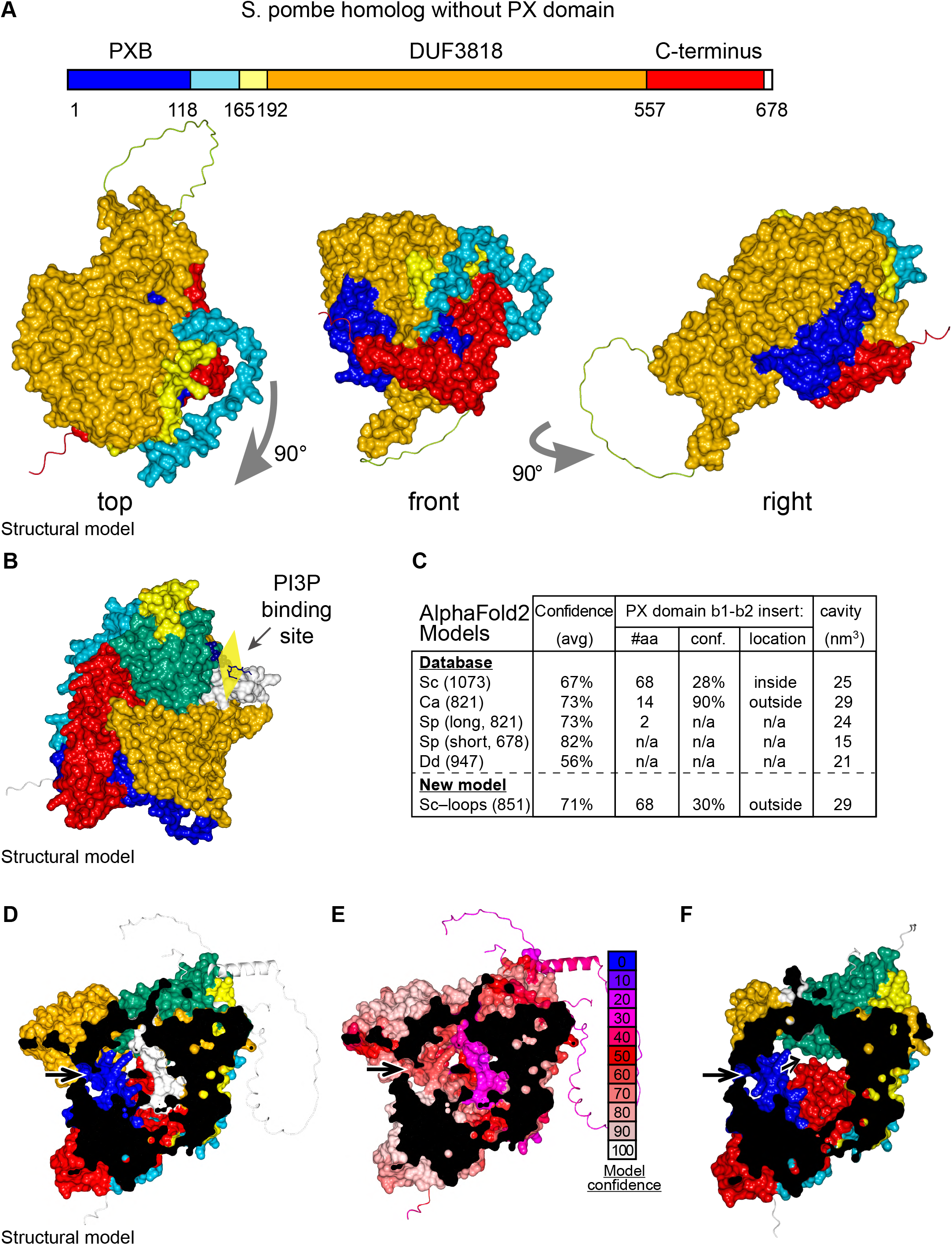
Details of AlphaFold2 structural predictions of Ypr097w in various species. (**A**) Domain map and surface representations of SPCC663.15c, the short homolog of Ypr097w in *S. pombe* lacking a PX domain. Coloring as in Fig. 4A-B. The domains are intimately intertwined, in particular the N-terminal PXB domain (blue) and extreme C-terminus (red). (**B**) Model of Ypr097w with super-posed headgroup of phosphatidylinositol 3-phosphate, by aligning the PX domain with that of p40phox (PDB 1H6H) (Bravo et al., 2001). (**C**) Key information about AlphaFold2 predictions for Ypr097w homologs. Data is from five pre- made AlphaFold2 models containing PXB and DUF3818: *S. cerevisiae* (Sc), *C. albicans* orf19.5621 (Ca), *S. pombe* SPCC1450.12 and SPCC663.15c (Sp 821 and 678 aa respectively) and *D. discoideum* (Dd) Q54JB8 (Jumper et al., 2021), together with a new model of Ypr097w (851 aa, see part F). Data shown: number of residues, overall confidence across the whole model (average pLDDT for all residues), details for the insert between strand 1 and strand 2 of the PX domain where *S. cerevisiae* Ypr097w uniquely has a loop with a local confidence score well below the 50% cut-off for useful interpretation (AlQuraishi, 2021), and estimated size of the internal cavity as calculated by MOLE. **(D-E)** AlphaFold2 prediction for the whole of Ypr097w, clipped to reveal the internal cavity, coloring either by domain (D, as in Fig. 4A-B), or by conservation (E, as in the key). Residues 243-280 in the first loop run internally (white in D, purple in E) **(F)** AlphaFold2 prediction for the Ypr097w missing two loops (residues 305-322, 356-468) and one globular region (690-780) (see Methods). Note the white region adopts a new position (compare to D). In parts D-F, arrows indicate potential tunnels linking the cavity to the cytosol.

**Figure 5 S1.**
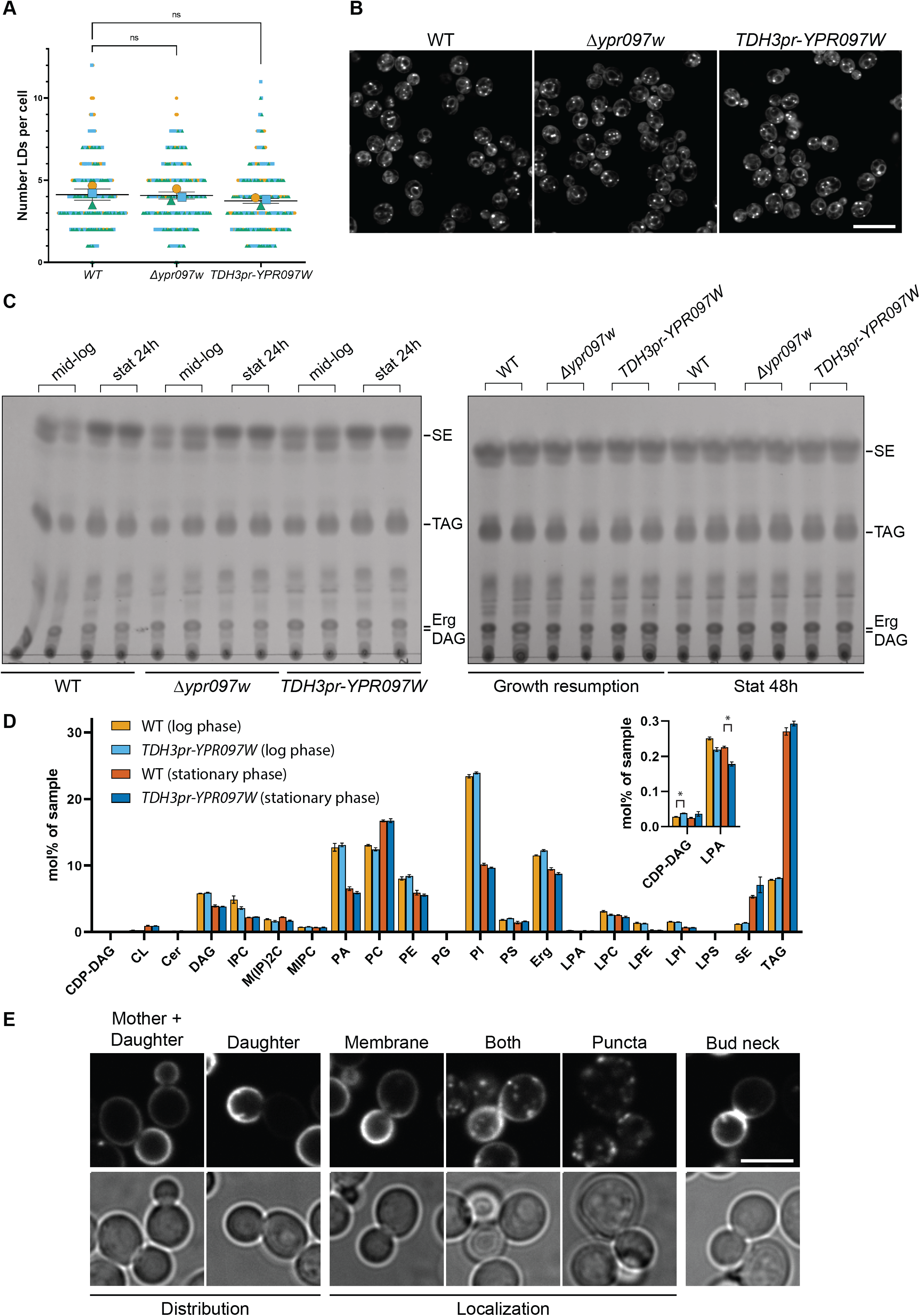
Ypr097w does not dramatically impact the total level of cellular lipids. (A) Quantification of LD number in WT, *Δypr097w* and overexpression *TDH3pr-YPR097W* cells, using the LD marker BODIPY. SuperPlot representation of individual cells (small symbols) and respective means (large symbols), from 3 independent experiments (different colors). Data are presented as mean ± SEM (n=3). One-way ANOVA with Dunnett’s multiple comparison test shows no significant differences between strains. ns. not significant. (B) LD morphology and distribution in WT, *Δypr097w* and overexpression *TDH3pr-YPR097W* cells, using the LD marker BODIPY. Scale bar, 10 µm; Images obtained using Micro 2. (C) Thin layer chromatography of WT, *Δypr097w* and overexpression *TDH3pr-YPR097W* cells in mid-logarithmic growth, stationary 24 h (left membrane), stationary 48 h and growth resumption (right membrane). SE, sterol esters; TAG, triacylglycerol; Erg, ergosterol; DAG, diacylglycerol. (D) Lipidomics analysis of WT and overexpression *TDH3pr-YPR097W* cells in mid-logarithmic growth and stationary 48 h. Acronyms as in (C) and CDP-DAG, cytidine diphosphate diacylglycerol; CL, cardiolipin; Cer, ceramide; IPC, inositolphosphorylceramide, M(IP)2C, mannosyl-di- (inositolphosphoryl) ceramide; MIPC, mannosyl-inositolphosphorylceramide; PA, phosphatidic acid; PC, phosphatidylcholine; PE, phosphatidylethanolamine; PG, phosphatidylglycerol; PI, phosphatidylinositol; PS, phosphatidylserine; LPA, lyso-phosphatidate; LPC, lyso-phosphatidylcholine; LPE, lyso-phosphatidylethanolamine; LPI, lyso- phosphatidylinositol; LPS, lyso-phosphatidylserine. (E) Examples of phenotypes used to quantify mCherry-D4H cellular distribution and localization. Scale bar, 5 µm; Images obtained using Micro 2.

## References

1. Alford SC, Ding Y, Simmen T, Campbell RE. 2012. Dimerization-Dependent Green and Yellow Fluorescent Proteins. ACS Synthetic Biology 1:569–575. doi:10.1021/SB300050J

2. AlQuraishi M. 2021. Protein-structure prediction revolutionized. Nature 2021 596:7873 **596**:487–488. doi:10.1038/d41586-021-02265-4

3. Anderson TM, Clay MC, Cioffi AG, Diaz KA, Hisao GS, Tuttle MD, Nieuwkoop AJ, Comellas G, Maryum N, Wang S, Uno BE, Wildeman EL, Gonen T, Rienstra CM, Burke MD. 2014. Amphotericin forms an extramembranous and fungicidal sterol sponge. Nature Chemical Biology 2014 10:5 10:400–406. doi:10.1038/nchembio.1496

4. Ashburner M, Ball CA, Blake JA, Botstein D, Butler H, Cherry JM, Davis AP, Dolinski K, Dwight SS, Eppig JT, Harris MA, Hill DP, Issel-Tarver L, Kasarskis A, Lewis S, Matese JC, Richardson JE, Ringwald M, Rubin GM, Sherlock G. 2000. Gene Ontology: tool for the unification of biology. Nature Genetics 2000 25:1 25:25–29. doi:10.1038/75556

5. Athenstaedt K. 2010. Isolation and Characterization of Lipid Particles from Yeast. Handbook of Hydrocarbon and Lipid Microbiology 4223–4229. doi:10.1007/978-3-540-77587-4_330

6. Bean BDM, Dziurdzik SK, Kolehmainen KL, Fowler CMS, Kwong WK, Grad LI, Davey M, Schluter C, Conibear E. 2018. Competitive organelle-specific adaptors recruit Vps13 to membrane contact sites. Journal of Cell Biology 217:3593–3607. doi:10.1083/JCB.201804111

7. Beltrao P, Albanèse V, Kenner LR, Swaney DL, Burlingame A, Villén J, Lim WA, Fraser JS, Frydman J, Krogan NJ. 2012. Systematic Functional Prioritization of Protein Posttranslational Modifications. Cell 150:413–425. doi:10.1016/J.CELL.2012.05.036

8. Brachmann CB, Davies A, Cost GJ, Caputo E, Li J, Hieter P, Boeke JD. 1998. Designer deletion strains derived fromSaccharomyces cerevisiae S288C: A useful set of strains and plasmids for PCR-mediated gene disruption and other applications. Yeast 14:115–132. doi:10.1002/(SICI)1097-0061(19980130)14:2<115::AID-YEA204>3.0.CO;2-2

9. Bravo J, Karathanassis D, Pacold CM, Pacold ME, Ellson CD, Anderson KE, Butler PJG, Lavenir I, Perisic O, Hawkins PT, Stephens L, Williams RL. 2001. The Crystal Structure of the PX Domain from p40phox Bound to Phosphatidylinositol 3-Phosphate. Molecular Cell 8:829–839. doi:10.1016/S1097-2765(01)00372-0

10. Castro IG, Schuldiner M, Zalckvar E. 2018. Mind the Organelle Gap – Peroxisome Contact Sites in Disease. Trends in Biochemical Sciences. doi:10.1016/j.tibs.2018.01.001

11. Chandra M, Chin YK-Y, Mas C, Feathers JR, Paul B, Datta S, Chen K-E, Jia X, Yang Z, Norwood SJ, Mohanty B, Bugarcic A, Teasdale RD, Henne WM, Mobli M, Collins BM. 2019. Classification of the human phox homology (PX) domains based on their phosphoinositide binding specificities. Nature Communications 2019 10:1 10:1–14. doi:10.1038/s41467-019-09355-y

12. Cho I-T, Adelmant G, Lim Y, Marto JA, Cho G, Golden JA. 2017. Ascorbate peroxidase proximity labeling coupled with biochemical fractionation identifies promoters of endoplasmic reticulum mitochondrial contacts. Journal of Biological Chemistry jbc.M117.795286. doi:10.1074/jbc.M117.795286

13. Cho KF, Branon TC, Rajeev S, Svinkina T, Udeshi ND, Thoudam T, Kwak C, Rhee H-W, Lee I- K, Carr SA, Ting AY. 2020. Split-TurboID enables contact-dependent proximity labeling in cells. Proceedings of the National Academy of Sciences 117:12143–12154. doi:10.1073/PNAS.1919528117

14. Chwastyk M, Panek EA, Malinowski J, Jaskólski M, Cieplak M. 2020. Properties of Cavities in Biological Structures—A Survey of the Protein Data Bank. Frontiers in Molecular Biosciences 0:314. doi:10.3389/FMOLB.2020.591381

15. Cieri D, Vicario M, Giacomello M, Vallese F, Filadi R, Wagner T, Pozzan T, Pizzo P, Scorrano L, Brini M, Calì T. 2018. SPLICS: A split green fluorescent protein-based contact site sensor for narrow and wide heterotypic organelle juxtaposition. Cell Death and Differentiation. doi:10.1038/s41418-017-0033-z

16. Cohen Y, Schuldiner M. 2011. Advanced Methods for High-Throughput Microscopy Screening of Genetically Modified Yeast Libraries. Methods in Molecular Biology 781:127–159. doi:10.1007/978-1-61779-276-2_8

17. Craene J-O de, Coleman J, Martin PE de, Pypaert M, Anderson S, John R., Yates I, Ferro-Novick S, Novick P. 2006. Rtn1p Is Involved in Structuring the Cortical Endoplasmic Reticulum. https://doi.org/101091/mbc.e06-01-0080 17:3009–3020. doi:10.1091/MBC.E06-01-0080

18. Currie E, Guo X, Christiano R, Chitraju C, Kory N, Harrison K, Haas J, Walther TC, Farese R v. 2014. High confidence proteomic analysis of yeast LDs identifies additional droplet proteins and reveals connections to dolichol synthesis and sterol acetylation [S]. Journal of Lipid Research 55:1465–1477. doi:10.1194/JLR.M050229

19. Daum G, Bohni PC, Schatz G. 1982. Import of proteins into mitochondria. Cytochrome b2 and cytochrome c peroxidase are located in the intermembrane space of yeast mitochondria. Journal of Biological Chemistry 257:13028–13033. doi:10.1016/S0021-9258(18)33617-2

20. Eisenberg-Bord M, Shai N, Schuldiner M, Bohnert M. 2016. A Tether Is a Tether Is a Tether: Tethering at Membrane Contact Sites. Developmental Cell 39:395–409. doi:10.1016/j.devcel.2016.10.022

21. Ejsing CS, Sampaio JL, Surendranath V, Duchoslav E, Ekroos K, Klemm RW, Simons K, Shevchenko A. 2009. Global analysis of the yeast lipidome by quantitative shotgun mass spectrometry. Proceedings of the National Academy of Sciences 106:2136–2141. doi:10.1073/PNAS.0811700106

22. Elbaz-Alon Y, Eisenberg-Bord M, Shinder V, Stiller SB, Shimoni E, Wiedemann N, Geiger T, Schuldiner M. 2015. Lam6 Regulates the Extent of Contacts between Organelles. Cell Reports 12:7–14. doi:10.1016/j.celrep.2015.06.022

23. Gabler F, Nam S-Z, Till S, Mirdita M, Steinegger M, Söding J, Lupas AN, Alva V. 2020. Protein Sequence Analysis Using the MPI Bioinformatics Toolkit. Current Protocols in Bioinformatics 72:e108. doi:10.1002/CPBI.108

24. Ganesan S, Tavassoli M, Shabits BN, Zaremberg V. 2020. Tubular ER Associates With Diacylglycerol-Rich Structures During Lipid Droplet Consumption. Frontiers in Cell and Developmental Biology 0:700. doi:10.3389/FCELL.2020.00700

25. Gatta AT, Wong LH, Sere YY, Calderón-Noreña DM, Cockcroft S, Menon AK, Levine TP. 2015. A new family of StART domain proteins at membrane contact sites has a role in ER-PM sterol transport. eLife 4:1–46. doi:10.7554/ELIFE.07253

26. Gietz RD, Woods RA. 2006. Yeast Transformation by the LiAc/SS Carrier DNA/PEG MethodYeast Protocols. New Jersey: Humana Press. pp. 107–120. doi:10.1385/1-59259-958-3:107

27. Gray KC, Palacios DS, Dailey I, Endo MM, Uno BE, Wilcock BC, Burke MD. 2012. Amphotericin primarily kills yeast by simply binding ergosterol. Proceedings of the National Academy of Sciences 109:2234–2239. doi:10.1073/PNAS.1117280109

28. Gueneau L, Fish RJ, Shamseldin HE, Voisin N, Tran Mau-Them F, Preiksaitiene E, Monroe GR, Lai A, Putoux A, Allias F, Ambusaidi Q, Ambrozaityte L, Cimbalistienė L, Delafontaine J, Guex N, Hashem M, Kurdi W, Jamuar SS, Ying LJ, Bonnard C, Pippucci T, Pradervand S, Roechert B, van Hasselt PM, Wiederkehr M, Wright CF, Xenarios I, van Haaften G, Shaw- Smith C, Schindewolf EM, Neerman-Arbez M, Sanlaville D, Lesca G, Guibaud L, Reversade B, Chelly J, Kučinskas V, Alkuraya FS, Reymond A. 2018. KIAA1109 Variants Are Associated with a Severe Disorder of Brain Development and Arthrogryposis. American Journal of Human Genetics 102:116–132. doi:10.1016/j.ajhg.2017.12.002

29. Hariri H, Speer N, Bowerman J, Rogers S, Fu G, Reetz E, Datta S, Feathers JR, Ugrankar R, Nicastro D, Henne WM. 2019. Mdm1 maintains endoplasmic reticulum homeostasis by spatially regulating lipid droplet biogenesis. Journal of Cell Biology 218:1319–1334. doi:10.1083/JCB.201808119

30. Henne WM, Zhu L, Balogi Z, Stefan C, Pleiss JA, Emr SD. 2015. Mdm1/Snx13 is a novel ER– endolysosomal interorganelle tethering protein. Journal of Cell Biology 210:541–551. doi:10.1083/JCB.201503088

31. Herker E, Vieyres G, Beller M, Krahmer N, Bohnert M. 2021. Lipid Droplet Contact Sites in Health and Disease. Trends in Cell Biology 31:345–358. doi:10.1016/J.TCB.2021.01.004

32. Herzog R, Schuhmann K, Schwudke D, Sampaio JL, Bornstein SR, Schroeder M, Shevchenko A. 2012. LipidXplorer: A Software for Consensual Cross-Platform Lipidomics. PLOS ONE 7:e29851. doi:10.1371/JOURNAL.PONE.0029851

33. Herzog R, Schwudke D, Schuhmann K, Sampaio JL, Bornstein SR, Schroeder M, Shevchenko A. 2011. A novel informatics concept for high-throughput shotgun lipidomics based on the molecular fragmentation query language. Genome Biology 2011 12:1 12:1–25. doi:10.1186/GB-2011-12-1-R8

34. Hugenroth M, Bohnert M. 2020. Come a little bit closer! Lipid droplet-ER contact sites are getting crowded. Biochimica et Biophysica Acta (BBA) - Molecular Cell Research 1867:118603. doi:10.1016/J.BBAMCR.2019.118603

35. Huh W-K, Falvo J v., Gerke LC, Carroll AS, Howson RW, Weissman JS, O’Shea EK. 2003. Global analysis of protein localization in budding yeast. Nature 2003 425:6959 425:686–691. doi:10.1038/nature02026

36. Hu J, Shibata Y, Zhu P-P, Voss C, Rismanchi N, Prinz WA, Rapoport TA, Blackstone C. 2009. A Class of Dynamin-like GTPases Involved in the Generation of the Tubular ER Network. Cell 138:549–561. doi:10.1016/J.CELL.2009.05.025

37. Johnson BB, Moe PC, Wang D, Rossi K, Trigatti BL, Heuck AP. 2012. Modifications in Perfringolysin O Domain 4 Alter the Cholesterol Concentration Threshold Required for Binding. Biochemistry 51:3373–3382. doi:10.1021/BI3003132

38. Jumper J, Evans R, Pritzel A, Green T, Figurnov M, Ronneberger O, Tunyasuvunakool K, Bates R, Žídek A, Potapenko A, Bridgland A, Meyer C, Kohl SAA, Ballard AJ, Cowie A, Romera- Paredes B, Nikolov S, Jain R, Adler J, Back T, Petersen S, Reiman D, Clancy E, Zielinski M, Steinegger M, Pacholska M, Berghammer T, Bodenstein S, Silver D, Vinyals O, Senior AW, Kavukcuoglu K, Kohli P, Hassabis D. 2021. Highly accurate protein structure prediction with AlphaFold. Nature 2021 596:7873 596:583–589. doi:10.1038/s41586-021-03819-2

39. Kakimoto Y, Tashiro S, Kojima R, Morozumi Y, Endo T, Tamura Y. 2018. Visualizing multiple inter-organelle contact sites using the organelle-targeted split-GFP system. Scientific Reports 2018 8:1 8:1–13. doi:10.1038/s41598-018-24466-0

40. Kane MS, Diamonstein CJ, Hauser N, Deeken JF, Niederhuber JE, Vilboux T. 2019. Endosomal trafficking defects in patient cells with KIAA1109 biallelic variants. Genes and Diseases. doi:10.1016/j.gendis.2018.12.004

41. Kelley LA, Mezulis S, Yates CM, Wass MN, Sternberg MJE. 2015. The Phyre2 web portal for protein modeling, prediction and analysis. Nature Protocols 2015 10:6 10:845–858. doi:10.1038/nprot.2015.053

42. Kishimoto T, Mioka T, Itoh E, Williams DE, Andersen RJ, Tanaka K. 2021. Phospholipid flippases and Sfk1 are essential for the retention of ergosterol in the plasma membrane. https://doi.org/101091/mbcE20-11-0699 mbc.E20-11-0699. doi:10.1091/MBC.E20-11-0699

43. Klose C, Surma MA, Gerl MJ, Meyenhofer F, Shevchenko A, Simons K. 2012. Flexibility of a Eukaryotic Lipidome – Insights from Yeast Lipidomics. PLOS ONE 7:e35063. doi:10.1371/JOURNAL.PONE.0035063

44. Kumagai K, Kawano-Kawada M, Hanada K. 2014. Phosphoregulation of the Ceramide Transport Protein CERT at Serine 315 in the Interaction with VAMP-associated Protein (VAP) for Inter-organelle Trafficking of Ceramide in Mammalian Cells *. Journal of Biological Chemistry 289:10748–10760. doi:10.1074/JBC.M113.528380

45. Kumar N, Leonzino M, Hancock-Cerutti W, Horenkamp FA, Li P, Lees JA, Wheeler H, Reinisch KM, de Camilli P. 2018. VPS13A and VPS13C are lipid transport proteins differentially localized at ER contact sites. Journal of Cell Biology 217:3625–3639. doi:10.1083/JCB.201807019

46. Kwak C, Shin S, Park J-S, Jung M, Nhung TTM, Kang M-G, Lee C, Kwon T-H, Park SK, Mun JY, Kim J-S, Rhee H-W. 2020. Contact-ID, a tool for profiling organelle contact sites, reveals regulatory proteins of mitochondrial-associated membrane formation. Proceedings of the National Academy of Sciences 117:12109–12120. doi:10.1073/PNAS.1916584117

47. Ladner CL, Yang J, Turner RJ, Edwards RA. 2004. Visible fluorescent detection of proteins in polyacrylamide gels without staining. Analytical Biochemistry 326:13–20. doi:10.1016/J.AB.2003.10.047

48. Laemmli UK. 1970. Cleavage of Structural Proteins during the Assembly of the Head of Bacteriophage T4. Nature 1970 227:5259 227:680–685. doi:10.1038/227680a0

49. Lamprecht MR, Sabatini DM, Carpenter AE. 2007. CellProfiler^TM^: free, versatile software for automated biological image analysis. https://doi.org/102144/000112257 42:71–75. doi:10.2144/000112257

50. Lang AB, John Peter ATAT, Walter P, Kornmann B. 2015. ER-mitochondrial junctions can be bypassed by dominant mutations in the endosomal protein Vps13. Journal of Cell Biology 210:883–890. doi:10.1083/jcb.201502105

51. Lanz MC, Yugandhar K, Gupta S, Sanford EJ, Faça VM, Vega S, Joiner AMN, Fromme JC, Yu H, Smolka MB. 2021. In-depth and 3-dimensional exploration of the budding yeast phosphoproteome. EMBO reports 22:e51121. doi:10.15252/EMBR.202051121

52. Levine TP, Munro S. 2001. Dual Targeting of Osh1p, a Yeast Homologue of Oxysterol-binding Protein, to both the Golgi and the Nucleus-Vacuole Junction. https://doi.org/101091/mbc1261633 12:1633–1644. doi:10.1091/MBC.12.6.1633

53. Liebisch G, Binder M, Schifferer R, Langmann T, Schulz B, Schmitz G. 2006. High throughput quantification of cholesterol and cholesteryl ester by electrospray ionization tandem mass spectrometry (ESI-MS/MS). Biochimica et Biophysica Acta (BBA) - Molecular and Cell Biology of Lipids 1761:121–128. doi:10.1016/J.BBALIP.2005.12.007

54. Li P, Lees JA, Lusk CP, Reinisch KM. 2020. Cryo-EM reconstruction of a VPS13 fragment reveals a long groove to channel lipids between membranes. Journal of Cell Biology 219. doi:10.1083/JCB.202001161

55. Liu T, Zhang XY, He XH, Geng JS, Liu Y, Kong DJ, Shi QY, Liu F, Wei W, Pang D. 2014. High levels of BCOX1 expression are associated with poor prognosis in patients with invasive ductal carcinomas of the breast. PLoS ONE 9:1–8. doi:10.1371/journal.pone.0086952

56. Loewen CJR, Roy A, Levine TP. 2003. A conserved ER targeting motif in three families of lipid binding proteins and in Opi1p binds VAP. The EMBO Journal 22:2025–2035. doi:10.1093/EMBOJ/CDG201

57. Maeda S, Otomo C, Otomo T. 2019. The autophagic membrane tether ATG2A transfers lipids between membranes. eLife 8. doi:10.7554/ELIFE.45777

58. Maekawa M, Fairn GD. 2015. Complementary probes reveal that phosphatidylserine is required for the proper transbilayer distribution of cholesterol. Journal of Cell Science 128:1422– 1433. doi:10.1242/JCS.164715

59. Marek M, Vincenzetti V, Martin SG. 2020. Sterol biosensor reveals LAM-family Ltc1-dependent sterol flow to endosomes upon Arp2/3 inhibition. Journal of Cell Biology 219. doi:10.1083/JCB.202001147

60. Mattia T di, Martinet A, Ikhlef S, McEwen AG, Nominé Y, Wendling C, Poussin-Courmontagne P, Voilquin L, Eberling P, Ruffenach F, Cavarelli J, Slee J, Levine TP, Drin G, Tomasetto C, Alpy F. 2020. FFAT motif phosphorylation controls formation and lipid transfer function of inter-organelle contacts. The EMBO Journal 39:e104369. doi:10.15252/EMBJ.2019104369

61. Mi H, Muruganujan A, Ebert D, Huang X, Thomas PD. 2019. PANTHER version 14: more genomes, a new PANTHER GO-slim and improvements in enrichment analysis tools. Nucleic Acids Research 47:D419–D426. doi:10.1093/NAR/GKY1038

62. Mirdita M, Ovchinnikov S, Steinegger M. 2021. ColabFold - Making protein folding accessible to all. bioRxiv 2021.08.15.456425. doi:10.1101/2021.08.15.456425

63. Mistry J, Chuguransky S, Williams L, Qureshi M, Salazar GA, Sonnhammer ELL, Tosatto SCE, Paladin L, Raj S, Richardson LJ, Finn RD, Bateman A. 2021. Pfam: The protein families database in 2021. Nucleic Acids Research 49:D412–D419. doi:10.1093/NAR/GKAA913

64. Mueller GA, Pedersen LC, Lih FB, Glesner J, Moon AF, Chapman MD, Tomer KB, London RE, Pomés A. 2013. The novel structure of the cockroach allergen Bla g 1 has implications for allergenicity and exposure assessment. Journal of Allergy and Clinical Immunology 132:1420–1426.e9. doi:10.1016/J.JACI.2013.06.014

65. Murley A, Sarsam RD, Toulmay A, Yamada J, Prinz WA, Nunnari J. 2015. Ltc1 is an ER- localized sterol transporter and a component of ER–mitochondria and ER–vacuole contacts. Journal of Cell Biology 209:539–548. doi:10.1083/JCB.201502033

66. Murphy SE, Levine TP. 2016. VAP, a Versatile Access Point for the Endoplasmic Reticulum: Review and analysis of FFAT-like motifs in the VAPome. Biochimica et Biophysica Acta - Molecular and Cell Biology of Lipids 1861:952–961. doi:10.1016/j.bbalip.2016.02.009

67. Neuman SD, Bashirullah A. 2018. Hobbit regulates intracellular trafficking to drive insulin- dependent growth during Drosophila development. Development 145:dev161356. doi:10.1242/dev.161356

68. Neuman SD, Jorgensen JR, Cavanagh AT, Smyth JT, Selegue JE, Emr SD, Bashirullah A. 2022. The Hob proteins are novel and conserved lipid-binding proteins at ER–PM contact sites. Journal of Cell Science 135. doi:10.1242/JCS.259086

69. Osawa T, Kotani T, Kawaoka T, Hirata E, Suzuki K, Nakatogawa H, Ohsumi Y, Noda NN. 2019. Atg2 mediates direct lipid transfer between membranes for autophagosome formation. Nature Structural & Molecular Biology 2019 26:4 26:281–288. doi:10.1038/s41594-019-0203-4

70. Pan X, Roberts P, Chen Y, Kvam E, Shulga N, Huang K, Lemmon S, Goldfarb DS. 2000. Nucleus–Vacuole Junctions in Saccharomyces cerevisiae Are Formed Through the Direct Interaction of Vac8p with Nvj1p. https://doi.org/101091/mbc1172445 11:2445–2457. doi:10.1091/MBC.11.7.2445

71. Park J-S, Thorsness MK, Policastro R, McGoldrick LL, Hollingsworth NM, Thorsness PE, Neiman AM. 2016. Yeast Vps13 promotes mitochondrial function and is localized at membrane contact sites. Molecular Biology of the Cell 27:2435–2449. doi:10.1091/mbc.E16-02-0112

72. Peter ATJ, van Schie SNS, Cheung NJ, Michel AH, Peter M, Kornmann B. 2021. Rewiring phospholipid biosynthesis reveals robustness in membrane homeostasis and uncovers lipid regulatory players. bioRxiv 2021.07.20.453065. doi:10.1101/2021.07.20.453065

73. Pettersen EF, Goddard TD, Huang CC, Couch GS, Greenblatt DM, Meng EC, Ferrin TE. 2004. UCSF Chimera—A visualization system for exploratory research and analysis. Journal of Computational Chemistry 25:1605–1612. doi:10.1002/JCC.20084

74. Pettersen EF, Goddard TD, Huang CC, Meng EC, Couch GS, Croll TI, Morris JH, Ferrin TE. 2021. UCSF ChimeraX: Structure visualization for researchers, educators, and developers. Protein Science 30:70–82. doi:10.1002/PRO.3943

75. Pravda L, Sehnal D, Toušek D, Navrátilová V, Bazgier V, Berka K, Svobodová Vařeková R, Koča J, Otyepka M. 2018. MOLEonline: a web-based tool for analyzing channels, tunnels and pores (2018 update). Nucleic Acids Research 46:W368–W373. doi:10.1093/NAR/GKY309

76. Prinz WA, Toulmay A, Balla T. 2020. The functional universe of membrane contact sites. Nature Reviews Molecular Cell Biology. doi:10.1038/s41580-019-0180-9

77. Rogers S, Hariri H, Wood NE, Speer NO, Henne WM. 2021. Glucose restriction drives spatial reorganization of mevalonate metabolism. eLife 10. doi:10.7554/ELIFE.62591

78. Scorrano L, Matteis MA de, Emr S, Giordano F, Hajnóczky G, Kornmann B, Lackner LL, Levine TP, Pellegrini L, Reinisch K, Rizzuto R, Simmen T, Stenmark H, Ungermann C, Schuldiner M. 2019. Coming together to define membrane contact sites. Nature Communications 2019 10:1 10:1–11. doi:10.1038/s41467-019-09253-3

79. Shai N, Yifrach E, van Roermund CWT, Cohen N, Bibi C, Ijlst L, Cavellini L, Meurisse J, Schuster R, Zada L, Mari MC, Reggiori FM, Hughes AL, Escobar-Henriques M, Cohen MM, Waterham HR, Wanders RJA, Schuldiner M, Zalckvar E. 2018. Systematic mapping of contact sites reveals tethers and a function for the peroxisome-mitochondria contact. Nature Communications 9:1761-undefined. doi:10.1038/s41467-018-03957-8

80. Shin JJH, Liu P, Chan LJ, Stefan C, Smits GJ, Loewen CJR. 2020. pH Biosensing by PI4P Regulates Cargo Sorting at the TGN Article pH Biosensing by PI4P Regulates Cargo Sorting at the TGN. Developmental Cell 1–16. doi:10.1016/j.devcel.2019.12.010

81. Slee JA, Levine TP. 2019. Systematic Prediction of FFAT Motifs Across Eukaryote Proteomes Identifies Nucleolar and Eisosome Proteins With the Predicted Capacity to Form Bridges to the Endoplasmic Reticulum: https://doi.org/101177/2515256419883136 2:251525641988313. doi:10.1177/2515256419883136

82. Song J, Yang W, Shih IM, Zhang Z, Bai J. 2006. Identification of BCOX1, a novel gene overexpressed in breast cancer. Biochimica et Biophysica Acta - General Subjects 1760:62–69. doi:10.1016/j.bbagen.2005.09.017

83. Sung M-K, Huh W-K. 2007. Bimolecular fluorescence complementation analysis system for in vivo detection of protein–protein interaction in Saccharomyces cerevisiae. Yeast 24:767– 775. doi:10.1002/YEA.1504

84. Surma MA, Herzog R, Vasilj A, Klose C, Christinat N, Morin-Rivron D, Simons K, Masoodi M, Sampaio JL. 2015. An automated shotgun lipidomics platform for high throughput, comprehensive, and quantitative analysis of blood plasma intact lipids. European Journal of Lipid Science and Technology 117:1540–1549. doi:10.1002/EJLT.201500145

85. Tamura N, Nishimura T, Sakamaki Y, Koyama-Honda I, Yamamoto H, Mizushima N. 2017. Differential requirement for ATG2A domains for localization to autophagic membranes and lipid droplets. FEBS Letters 591:3819–3830. doi:10.1002/1873-3468.12901

86. Tang Z, Takahashi Y, He H, Hattori T, Chen C, Liang X, Chen H, Young MM, Wang H-G. 2019. TOM40 Targets Atg2 to Mitochondria-Associated ER Membranes for Phagophore Expansion. Cell Reports 28:1744–1757.e5. doi:10.1016/J.CELREP.2019.07.036

87. Tarassov K, Messier V, Landry CR, Radinovic S, Serna Molina MM, Shames I, Malitskaya Y, Vogel J, Bussey H, Michnick SW. 2008. An in vivo map of the yeast protein interactome. Science 320:1465–1470. doi:10.1126/SCIENCE.1153878

88. Tashiro S, Kakimoto Y, Shinmyo M, Fujimoto S, Tamura Y. 2020. Improved Split-GFP Systems for Visualizing Organelle Contact Sites in Yeast and Human Cells. Frontiers in Cell and Developmental Biology 8:1353. doi:10.3389/fcell.2020.571388

89. The Gene Ontology Consortium. 2019. The Gene Ontology Resource: 20 years and still GOing strong. Nucleic Acids Research 47:D330–D338. doi:10.1093/NAR/GKY1055

90. Tong AHY, Boone C. 2006. Synthetic Genetic Array Analysis in Saccharomyces cerevisiae. Methods in molecular biology (Clifton, NJ) 313:171–191. doi:10.1385/1-59259-958-3:171

91. Ugrankar R, Bowerman J, Hariri H, Chandra M, Chen K, Bossanyi MF, Datta S, Rogers S, Eckert KM, Vale G, Victoria A, Fresquez J, McDonald JG, Jean S, Collins BM, Henne WM. 2019. Drosophila Snazarus Regulates a Lipid Droplet Population at Plasma Membrane- Droplet Contacts in Adipocytes. Developmental Cell 50:557–572.e5. doi:10.1016/J.DEVCEL.2019.07.021

92. Valm AM, Cohen S, Legant WR, Melunis J, Hershberg U, Wait E, Cohen AR, Davidson MW, Betzig E, Lippincott-Schwartz J. 2017. Applying systems-level spectral imaging and analysis to reveal the organelle interactome. Nature 546:162–167. doi:10.1038/nature22369

93. Valverde DP, Yu S, Boggavarapu V, Kumar N, Lees JA, Walz T, Reinisch KM, Melia TJ. 2019. ATG2 transports lipids to promote autophagosome biogenesis. Journal of Cell Biology 218:1787–1798. doi:10.1083/JCB.201811139

94. van Vliet AR, Giordano F, Gerlo S, Segura I, van Eygen S, Molenberghs G, Rocha S, Houcine A, Derua R, Verfaillie T, Vangindertael J, de Keersmaecker H, Waelkens E, Tavernier J, Hofkens J, Annaert W, Carmeliet P, Samali A, Mizuno H, Agostinis P. 2017. The ER Stress Sensor PERK Coordinates ER-Plasma Membrane Contact Site Formation through Interaction with Filamin-A and F-Actin Remodeling. Molecular Cell 65:885–899.e6. doi:10.1016/j.molcel.2017.01.020

95. Velikkakath AKG, Nishimura T, Oita E, Ishihara N, Mizushima N. 2012. Mammalian Atg2 proteins are essential for autophagosome formation and important for regulation of size and distribution of lipid droplets. https://doi.org/101091/mbc.e11-09-0785 23:896–909. doi:10.1091/MBC.E11-09-0785

96. Voeltz GK, Prinz WA, Shibata Y, Rist JM, Rapoport TA. 2006. A Class of Membrane Proteins Shaping the Tubular Endoplasmic Reticulum. Cell 124:573–586. doi:10.1016/J.CELL.2005.11.047

97. Weill U, Yofe I, Sass E, Stynen B, Davidi D, Natarajan J, Ben-Menachem R, Avihou Z, Goldman O, Harpaz N, Chuartzman S, Kniazev K, Knoblach B, Laborenz J, Boos F, Kowarzyk J, Ben-Dor S, Zalckvar E, Herrmann JM, Rachubinski RA, Pines O, Rapaport D, Michnick SW, Levy ED, Schuldiner M. 2018. Genome-wide SWAp-Tag yeast libraries for proteome exploration. Nature Methods 2018 15:8 15:617–622. doi:10.1038/s41592-018-0044-9

98. Yang J, Anishchenko I, Park H, Peng Z, Ovchinnikov S, Baker D. 2020. Improved protein structure prediction using predicted interresidue orientations. Proceedings of the National Academy of Sciences 117:1496–1503. doi:10.1073/PNAS.1914677117

99. Yang Z, Zhao X, Xu J, Shang W, Tong C. 2018. A novel fluorescent reporter detects plastic remodeling of mitochondria–ER contact sites. Journal of Cell Science 131. doi:10.1242/JCS.208686

100. Yeshaw WM, van der Zwaag M, Pinto F, Lahaye LL, Faber AIE, Gómez-Sánchez R, Dolga AM, Poland C, Monaco AP, van IJzendoorn SCD, Grzeschik NA, Velayos-Baeza A, Sibon OCM. 2019. Human VPS13A is associated with multiple organelles and influences mitochondrial morphology and lipid droplet motility. eLife 8. doi:10.7554/ELIFE.43561

101. Yofe I, Schuldiner M. 2014. Primers-4-Yeast: a comprehensive web tool for planning primers for Saccharomyces cerevisiae. Yeast 31:77–80. doi:10.1002/YEA.2998

102. Yofe I, Weill U, Meurer M, Chuartzman S, Zalckvar E, Goldman O, Ben-Dor S, Schütze C, Wiedemann N, Knop M, Khmelinskii A, Schuldiner M. 2016. One library to make them all: streamlining the creation of yeast libraries via a SWAp-Tag strategy. Nature methods 13:371–8. doi:10.1038/nmeth.3795

103. Yu JW, Lemmon MA. 2001. All Phox Homology (PX) Domains from Saccharomyces cerevisiae Specifically Recognize Phosphatidylinositol 3-Phosphate *. Journal of Biological Chemistry 276:44179–44184. doi:10.1074/JBC.M108811200

104. Zaremberg V, McMaster CR. 2002. Differential Partitioning of Lipids Metabolized by Separate Yeast Glycerol-3-phosphate Acyltransferases Reveals That Phospholipase D Generation of Phosphatidic Acid Mediates Sensitivity to Choline-containing Lysolipids and Drugs *. Journal of Biological Chemistry 277:39035–39044. doi:10.1074/JBC.M207753200

105. Zung N, Schuldiner M. 2020. New horizons in mitochondrial contact site research. Biological Chemistry 401:793–809. doi:10.1515/HSZ-2020-0133

